# Viscoadaptation controls intracellular reaction rates in response to heat and energy availability

**DOI:** 10.1101/709717

**Authors:** Laura Persson, Vardhaan S. Ambati, Onn Brandman

## Abstract

Cells must precisely orchestrate thousands of reactions in both time and space. Yet reaction kinetics are highly dependent on uncontrollable environmental conditions such as temperature. Here, we report a novel mechanism by which budding yeast influence reaction rates through adjustment of intracellular viscosity. This “viscoadaptation” is achieved by production of two carbohydrates, trehalose and glycogen, which combine to create a more viscous cellular environment in which biomolecules retain solubility. We demonstrate that viscoadaptation functions as both an acute response to temperature increase as well as a homeostatic mechanism, allowing cells grown at temperatures spanning from 22°C to 40°C to maintain equivalent rates of intracellular diffusion and diffusion-controlled chemical reactions. Multiple conditions that lower ATP trigger viscoadaptation, suggesting that viscoadaptation may be a general cellular response to low energy. Viscoadaptation reveals viscosity to be a tunable property of cells through which they can regulate diffusion-controlled processes dynamically in response to a changing environment.

## Introduction

Warm-blooded organisms thrive in a wide range of environments by maintaining nearly constant internal temperatures. Yet many organisms that lack homeothermic regulation also tolerate large and unpredictable fluctuations in temperature. The budding yeast *S. cerevisiae*, for example, can grow and divide at temperatures that span 30°C or more (*1, 2*). Other organisms have adapted to more extreme conditions and can survive temperatures ranging from 80°C to 120°C(*3*). Such versatility is remarkable given the numerous thermodynamic challenges that temperature variation poses for a cell. Many cellular processes, such as protein folding, are dictated by small energetic differences between states and are therefore highly sensitive to temperature(*4*). In fact, much of the seminal work on heat adaptation has focused on protein folding and the role of chaperones in creating folding microenvironments that are buffered against large variations in the energetics of the system as a whole(*5*).

In addition to these *intra*molecular processes, temperature has a profound effect on *inter*molecular processes. In general, increasing the temperature of a liquid lowers its viscosity thereby increasing the diffusion rate of any constituent particles (*6*). As such, particle collisions become more frequent and diffusion-controlled reactions proceed more quickly (*6*). However, it is largely unknown how these effects manifest in live cells with highly complex and dynamic interaction networks that may be affected by temperature to different extents. The interconnectedness of cellular pathways also suggests that perturbation of reaction kinetics in even one pathway has the potential to impact multiple aspects of cellular functioning. The extent of these challenges and the cellular strategies that may mitigate them have been largely unexplored.

Seminal work in the 1970s revealed a homeostatic mechanism for controlling the fluidity of cell membranes in response to changing temperatures. Coined homeoviscous adaptation, cells respond to changes in temperature by adjusting the lipid content of their membranes to maintain their appropriate viscosity (*7*). In this way, the cell is able to control the rate of membrane reactions in a temperature-independent way (*8*). No such mechanism has been identified for cytosolic reactions. In the following work, we compared the rate of an exogenous model reaction *in vivo* and *in vitro* as a function of temperature. In doing so, we discovered a form of cellular adaptation that renders a reaction rate invariant to temperature change *in vivo*, despite strong temperature dependence *in vitro*. Termed “viscoadaptation”, this response allows the cell to maintain a near-constant intracellular viscosity despite changes in the temperature of the environment. As a result, diffusion rates and viscosity-controlled reaction rates are held invariant across at least 20° C. Viscoadaptation is both a homeostatic mechanism for growth at multiple temperatures as well as an acute response to a variety of environmental stressors.

### Cells homeostatically regulate the rate of an exogenous reaction in response to temperature changes

To determine the effect of temperature on reaction rates in live cells, we made use of an exogenous biotinylation reaction in which the BirA enzyme from *E. coli* is co-expressed with its biotin-accepting substrate (the “Avi” tag) fused to a “bait” protein in yeast (*9, 10*) (Figure 1A). We measured the progress of this reaction by a gel shift assay in which the biotinylated species runs more slowly than its non-biotinylated counterpart. After initiation of the reaction by the addition of biotin to the media, the ratio of the biotinylated species to the non-biotinylated species increases with time (Figure 1A). Unless otherwise specified, reactions described here occur between BirA and GFP-Avi.

**Figure 1.**
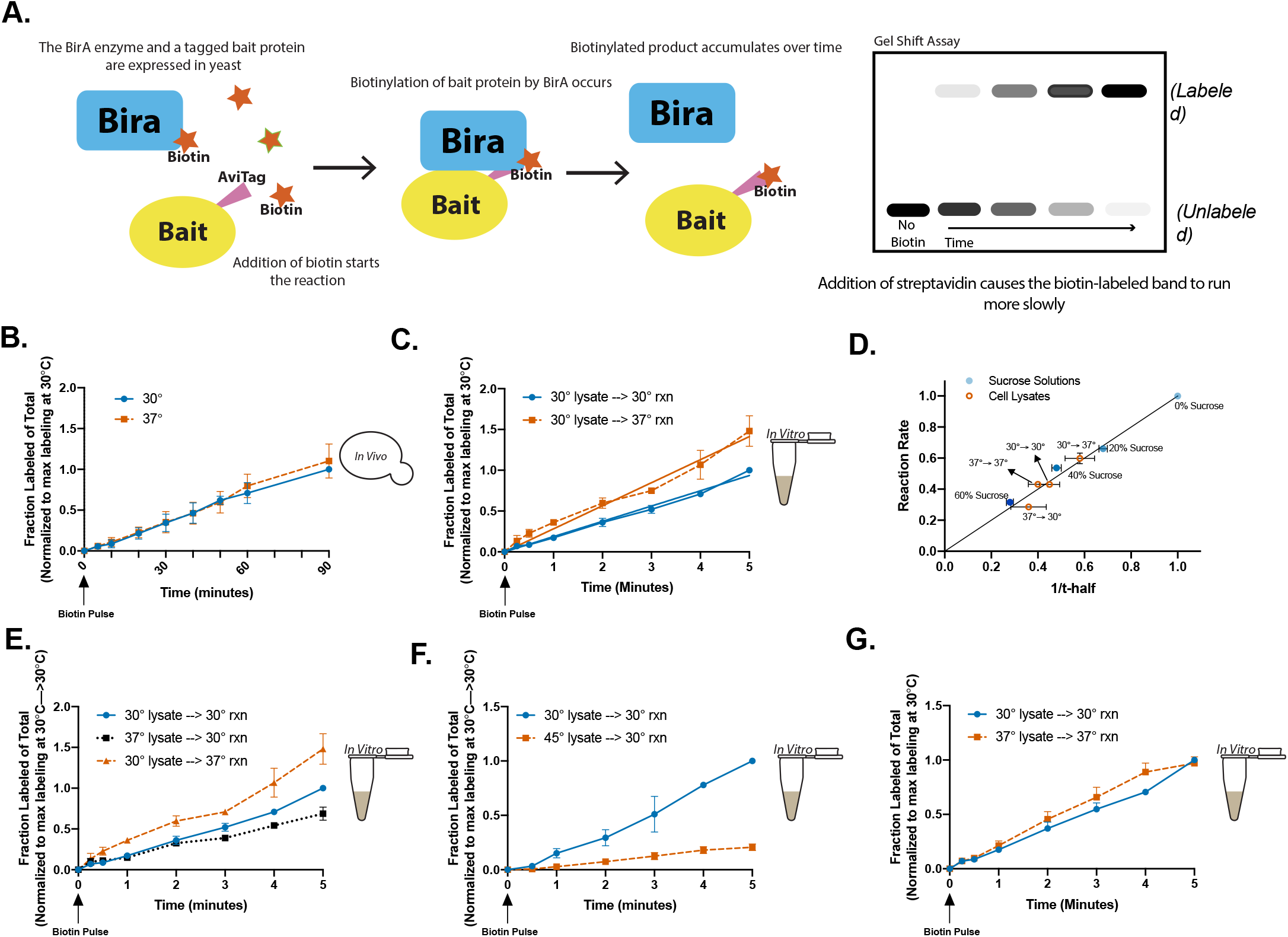
A viscosity-controlled model reaction is temperature dependent in cell lysates but not live cells. (**A**) BirA biotinylates a 15-AA target sequence called the Avi-tag. Labeling rates can be determined by addition of streptavidin which causes the biotinylated species to run at a higher molecular weight. All reactions use GFP-Avi as “bait”. (**B**) Biotinylation reaction rate in live yeast grown at 30°C (●blue) or 37°C (■orange). Normalized to max labeling at 30°C. (n= 4 time courses/temperature) (**C**) Reaction rates in cell lysates from unstressed cells at 30°C (●blue) or 37°C (■orange). All *in vitro* reactions are normalized to the max labeling for 30°C lysate at 30°C. (n= 5 time courses/condition) (**D**) Rate of GFP-Avi labeling vs. 1/t-half for reactions in 0%, 20%, 40%, or 60% sucrose solutions (●blue) or in cell lysates labeled as “growth temp. prior to lysis → measurement temp” (**○** orange). Rates and t-half values were normalized to their respective values in 0% sucrose (H_2_O only). Horizontal error bars are s.em. for ≤ 3 independent FRAP experiments. Vertical errror bars are calculated by plotting the best fit line for the average of multiple time courses (n ≥ 5 time courses/condition) and calculating the standard error of the slope. (**E**) *In vitro* reactions were carried out in the indicated lysates at the indicated temperatures. (n = 5 time courses/condition) (**F**) *In vitro* reactions performed at 30°C in lysates generated from 30°C cultures (●blue) or cultures subjected to 30 minutes at 45°C prior to lysis (■orange). (n= 3 time courses/condition) (**G**) *In vitro* reactions for which the reaction temperature is the same as the growth temperature of the cells prior to lysis. (n = 5 time courses/condition) Unless otherwise specified all error bars are s.e.m.

When we measured the rate of the BirA reaction in cells growing at 30°C or 37°C (normal growth temperatures for yeast), we found no difference in rate (Figure 1B). This *in vivo* temperature insensitivity could be caused by intrinsic properties of the reaction or depend on an active process in the cell that adapts to different temperatures. To explore this, we compared the behavior of the reaction *in vivo* to the behavior of the reaction in a cell-free system of undiluted yeast lysate. The reaction environment should be largely the same in lysates and cells, but any active cellular process that senses changes in the environment and initiates corresponding cellular adaptation should not be possible. To remove additional variables, equal amounts of exogenous BirA, ATP, and biotin were added to the lysates. We now observed a 50% increase in the rate of the reaction at 37°C relative to 30°C. The difference between *in vivo* and *in vitro* results suggests that live cells actively regulate the speed of the reaction (Figure 1C). Because the reaction is exogenous and highly specific to the *E. coli*-derived Avi sequence, there should be no evolved system in yeast for its specific regulation(*11*). We therefore predicted that these observations reflect a broader strategy of reaction regulation in cells.

We reasoned that the mechanism by which cells counteracted the effects of temperature would correspond to the mechanism by which temperature influences the biotinylation reaction rate. One possibility was that viscosity, a temperature-controlled property of the reaction medium, regulates the reaction rate by affecting molecular movement. To characterize the relationship between the reaction rate and viscosity we varied viscosity *in vitro* using different concentrations of sucrose (Figure 1D). Measuring solution viscosity by Fluorescence Recovery After Photobleaching (FRAP), we then plotted reaction rate versus viscosity (reported as 1/t-half). We found that the rate of the BirA reaction scaled 1:1 with solution viscosity for all concentrations of sucrose tested (0%, 20%, 40%, 60%) (Figure 1D). Furthermore, this relationship between viscosity and reaction rate held for cell lysates harvested and measured at different temperatures (Figure 1D). Finally, dilution of the lysate 1:50 with H_2_O resulted in an 80% decrease in viscosity and an 80% increase in the reaction rate (quantified as the ratio of product to substrate and therefore independent of substrate concentration) suggesting that the rate of the BirA reaction is viscosity-controlled under these conditions. (Supp. Figure 1A, 1B).

We hypothesized that a viscosity-increasing molecule (viscogen) produced in 37°C-grown cells could explain the difference between the *in vitro* and *in vivo* results. If so, the reaction should be slower in lysates from 37°C cultures compared to those from 30°C cultures. Consistent with this hypothesis, the reaction proceeded 25% slower in the 37°C-derived lysate (Figure 1E). The reaction was even slower (80%), in lysate derived from cells that had been more severely heat shocked (45°C for 30 minutes prior to lysis) suggesting that the mechanism operates to different extents depending on the degree or type of temperature difference (acute change vs. steady state growth conditions) (Figure 1F).

If cells possess a tunable mechanism for regulating viscosity, we hypothesized that cells growing in steady state at different temperatures might adapt homeostatically to maintain invariant viscosity. If so, we should find that the *in vitro* reaction rates are equivalent if the reactions are performed at the temperature at which the cells were grown prior to lysis (30° or 37°). Indeed, we found that changing the *in vitro* reaction temperature to match the growth temperature equalized the reaction rates (Figure 1G). To check that our findings were not specific to GFP-Avi, we performed similar experiments using RPL19b-Avi and observed consistent results. (Supp Figure 2A, 2B). We conclude from these observations that cells possess a mechanism to maintain the rate of a viscosity-controlled reaction at different growth temperatures.

### Cells regulate viscosity in response to temperature changes

We next directly tested whether cells regulate viscosity in response to temperature changes. Changes in intracellular viscosity should result in corresponding changes in the diffusion of molecules in the cytosol. We therefore quantified viscosity with Fluorescence Recovery After Photobleaching (FRAP) to measure the mobility of GFP *in vivo* and *in vitro* under different conditions. FRAP data is reported as t-half values, that is, the time required for a bleached spot to recover to half of its maximum fluorescence (Figure 2A). Thus, smaller t-half values indicate faster diffusion and larger values indicate slower diffusion (Figure 2A). To determine the effect of temperature on diffusion in the absence of active cellular regulation, we measured the recovery of GFP in cell lysate at 22°C, 30°C, 37°C, and 42°C. We observed that the recovery time was shorter at higher temperatures with a maximum decrease of 40% at 42°C relative to 22°C (Figure 2B). This result is consistent with the known role of temperature in regulating diffusion in solutions.

**Figure 2.**
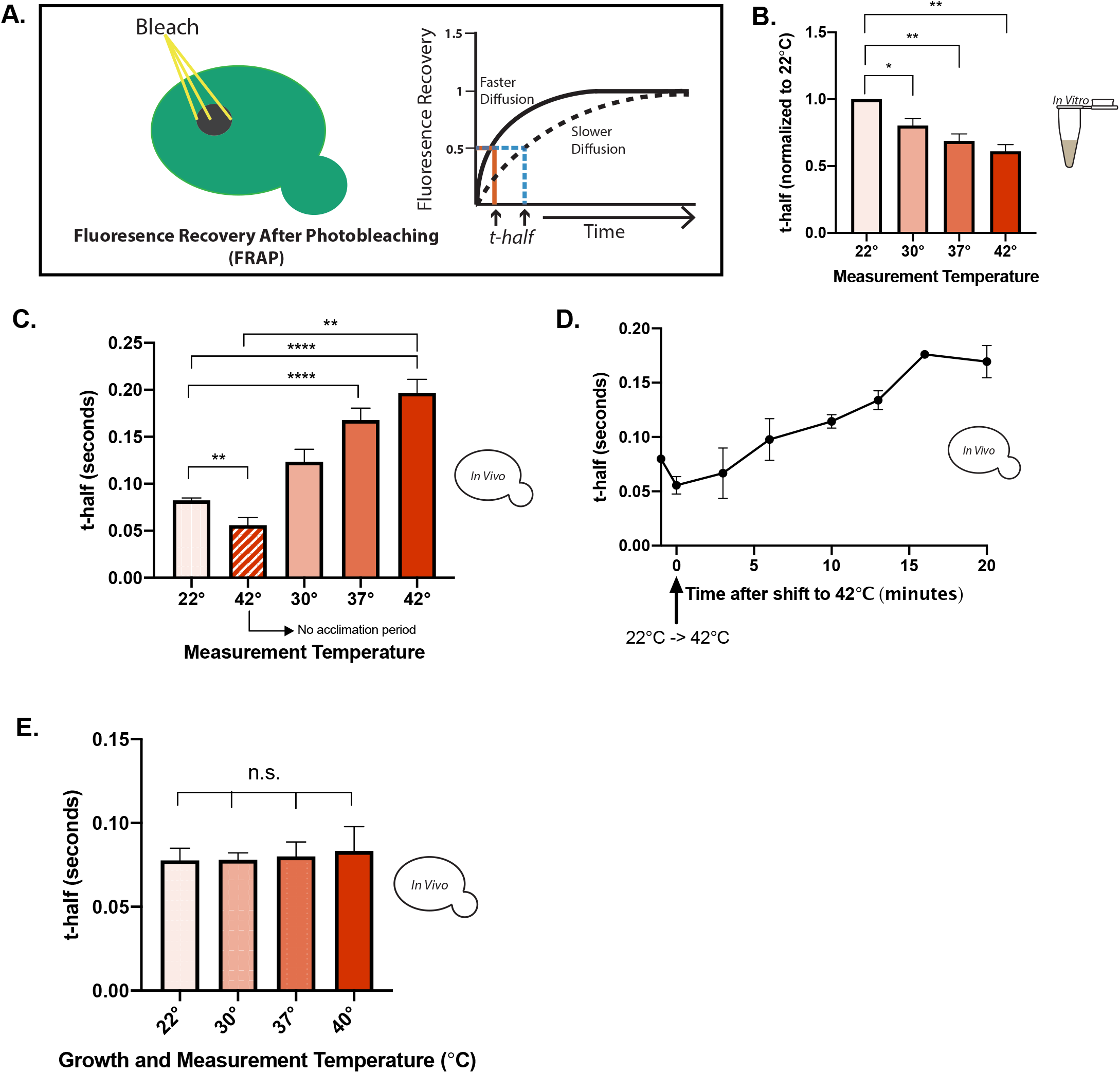
Cells regulate intracellular viscosity in response to heat (“viscoadaptation”). (**A**) Fluoresence Recovery After Photobleaching (FRAP) measurements on live yeast cells and cell lysates. Recovery is measured by t-half, the time to reach half-maximal fluoresence recovery. (**B**) Recovery of purified GFP in a cell lysate measured in a chamber heated to the indicated temperatures. Values are normalized to 22°C. (n= lysates/ temperature) (paired t-test 22°C vs 42°C p= .0015, 22°C vs 37°C p=.0041, 22°C vs 30°C p= .0198) (**C**) FRAP measurements in yeast grown at 22°C then given 20 minutes at the indicated temperatures before measurement. Striped bar represents cells measured immediately after the temperature change. (n≥ 3 experiments/temperature, >10 cells/experiment) (unpaired t-test 22°C vs 42°C p<.0001, 22°C vs 37°C p<.0001, immediate 42°C vs. delayed measurement at 42°C p=.0011)(**D**) FRAP measurements in live cells taken at the indicated times. First point is before temperature shift. T_0_ is the first time point after cells were transferred from 22°C to 42°C (n=4 timecourses) (**E**) Yeast cells grown to log phase at the indicated temperatures. FRAP measurments on GFP performed in a microscope chamber matched to the growth temperature of the cells. (n≥ 4 experiments, >10 cells/experiment) All error bars are s.e.m.

We next measured the diffusion of GFP in live cells across the same temperature differentials (22°C to 30°C, 37°C, or 42°C). Cells at 22°C had an average t-half of 0.08 seconds. When cells were transferred from 22°C to 42°C and measured within the first 5 minutes, the average t-half was 0.05 seconds, almost 40% lower and in agreement with the results in cell lysates. However, when cells were measured 20 minutes after the temperature shift, diffusion decreased with increasing temperature. In the most extreme case, the average t-half after 20 minutes of acclimation to 42°C was 0.19s, almost 300% higher than the t-half in cells measured immediately after the temperature shift (n ≥ 4 experiments, ≥15 cells per temperature/experiment) (Figure 2C). From these results, we conclude that cells respond to acute temperature increases by raising their viscosity, thereby slowing the diffusion of cytosolic proteins. We observed the same results using mKate2, a structurally dissimilar fluorophore, indicating that these observations were not unique to GFP (Supp Figure 3A, 3B).

To track how cells adapt over time to temperature increase, we collected a time course of FRAP measurements, beginning immediately after the temperature change (22°C to 42°C) and then every two minutes for 20 minutes (Figure 2D). Immediately after the temperature upshift, we saw a drop in recovery time to 0.05 seconds (~40% lower than the recovery time at 22), reflecting the effect of the temperature increase on diffusion prior to cellular adaptation. Yet by 5 minutes after the temperature change t-half values had begun to increase and did so gradually until plateauing around 0.18 seconds after 20 minutes, in agreement with prior results in cells measured 20 minutes after temperature shift. Because the mobility of GFP could be affected by other factors in the cell, we used an orthogonal method, microrheology with fluorescent beads, to measure the viscosity of cell lysates produced from cultures in different growth conditions. By FRAP and microrheology, we observed ~25% slower diffusion in lysates produced from heat shocked cells relative to unstressed cells (Supp Figure 4A,4B). However, during the course of these experiments, we observed the rapid formation of macroscopic, spherical inclusions selectively in lysate from heat shocked cells (Supp Figure 4C). These structures showed complete loss of interior mobility by microrheology (Supp Figure 4D). Measurements were therefore made in regions of the lysate that retained mobility, but as a result these values likely underestimate the viscosity of the lysate prior to inclusion formation. Based on these observations, we suggest that the phase properties of the cytosol in heat shocked cells may be highly unstable (addressed in more detail below).

Thus far, diffusion measurements have been made in cells experiencing acute heat stress. However, we observed homeostatic regulation of the BirA reaction at 30°C and 37°C. We therefore asked whether diffusion in the cytosol is also controlled homeostatically. To test this, we grew yeast to log-phase at 22°C, 30°C, 37°C, or 40°C and performed FRAP measurements at the growth temperature of the culture. Now, we found that the t-half for GFP was nearly constant at all temperatures, at a value of about 0.08 seconds (Figure 2E). Collectively, these results reveal that cells counteract the effect of temperature on protein diffusion by increasing their intracellular viscosity. We term this phenomenon “viscoadaptation”.

### Accumulation of trehalose and glycogen in response to heat underlies viscoadaptation

We next investigated the mechanism underlying viscoadaptation. We first tested whether this response was dependent on protein translation, as it is known that many specialized heat shock proteins are produced in response to temperature increase (*12, 13*). However, neither the degree nor speed of adaptation was significantly affected by the pretreatment with the translation inhibitor cycloheximide, indicating that protein synthesis is not required for viscoadaptation in response to acute heat shock (Supp Figure 5).

In addition to specialized heat shock proteins, temperature increase rapidly induces the synthesis of two carbohydrates, trehalose and glycogen(*14, 15*). Both trehalose and glycogen are assembled from glucose monomers and together can account for as much as 30% of a cell’s dry mass during severe heat shock(*16*) (Figure 3A). Trehalose and glycogen improve tolerance to heat stress, though the mechanisms underlying this function are not fully understood (*17, 18*). We adapted an iodine staining method to qualitatively determine changes in glycogen in live yeast and observed strong induction of glycogen in response to heat shock at 42°C (Figure 3C)(*19, 20*). A single glycogen molecule is composed of thousands of glucose monomers in an extensively branched spherical structure that can reach up to 40 nm in diameter in yeast(*21*). *In vitro*, glycogen acts as a viscogen(*22*).

**Figure 3.**
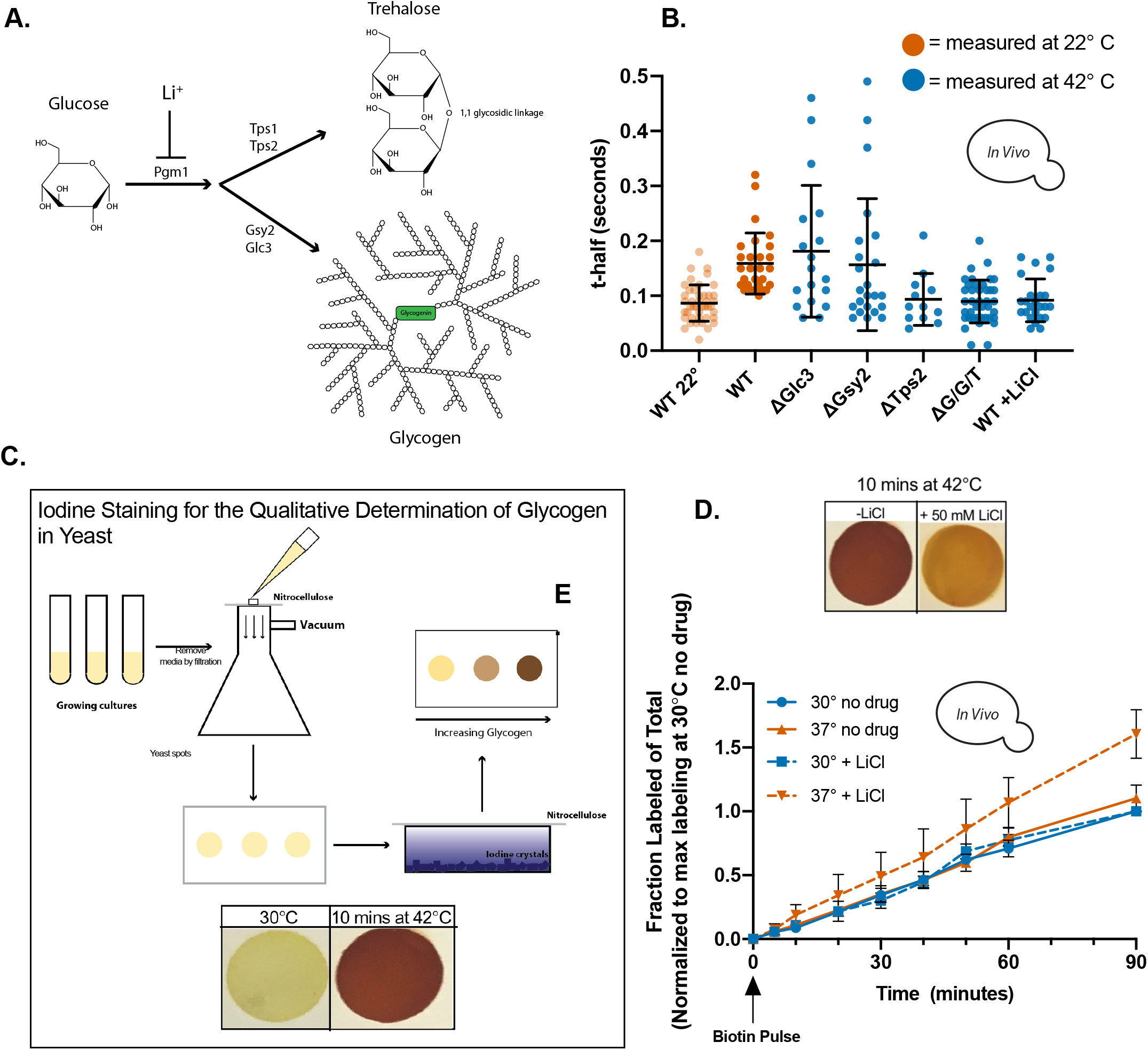
Heat-induced accumulation of the carbohydrates glycogen and trehalose is required for viscoadaptation. (**A**) Trehalose and glycogen are produced from glucose monomers. First, glucose-1-p is converted to glucose-6-p by the enzyme phosphoglucomutase (PGM). PGM activity can be inhibited by treatment with lithium. (**B**) GFP recovery times for the indicated strains and growth conditions. (**C**) Glycogen content in live yeast can be qualitatively assessed by staining with iodine vaper. 10 minutes of heat shock at 42°C causes accumulation of glycgen which can be abrogated by pretreatment with 50 mM LiCl (**D**) Cells expressing BirA and GFP-Avi were grown in steady state at 30°C (blue) or 37°C (orange) with or without the addition of 50 mM LiCl 1.5 hours prior to initiation of labeling (dashed and solid lines, respectively). All error bars are s.e.m.

In contrast to glycogen, trehalose is a small disaccharide formed by a 1,1-glycosidic bond between two α-glucose monomers(*23*). Trehalose has several unusual properties relevant to its biological function. According to the “immobilization theory”, trehalose protects against desiccation in cells by undergoing a liquid-to-glass phase transition that drastically reduces intracellular mobility(*23*). Trehalose has also been proposed to act as a kosmotrope, substituting for water around biomolecules and causing the surrounding water molecules to become more ordered(*23*). Both proposed functions reduce the mobility of biomolecules(*23*). We therefore asked how accumulation of trehalose and glycogen affects diffusion during heat stress. We first generated yeast mutants deficient in the synthesis of one or both carbohydrates. Yeast lacking the trehalose synthesis enzyme TPS2 were subjected to heat shock, and GFP diffusion was measured by FRAP. Whereas t-half values for the wildtype strain increased to 0.18s, t-half values for the ΔTPS2 strain remained the same (Figure 3B). This implies that trehalose production is necessary for viscoadaptation in response to heat shock.

We next examined diffusion after temperature increase in strains lacking the glycogen synthase, GSY2, or the glycogen branching enzyme, GLC3. Yeast deficient in glycogen synthesis demonstrated a wide range of phenotypes. Approximately 50% of cells fail to slow diffusion in response to heat (“non-responders”). However, the remainder of cells respond normally or hyper-respond (“responders”)(Figure 3B). Because trehalose and glycogen are made from the same precursor, disruption of one pathway can increase flux into the other (*24*) (Figure 3A). We therefore reasoned that the hyper-viscous sub-population of glycogen deficient cells could result from increased flux through the trehalose synthesis pathway and resultant over-accumulation of trehalose. Consistent with this model, when production of both trehalose and glycogen was impaired, all cells became non-responders (Figure 3B).

To determine how trehalose and glycogen affect reaction rates, we measured the rate of the BirA reaction in cells with compromised glycogen or trehalose production. However, we found that ΔTPS2 yeast could not grow in the low biotin media required for biotinylation assays. Glycogen deficient strains did not show a consistent effect on reaction rate, presumably due to the heterogeneity of the response at the single-cell level (Supp Figure 6). We next used a pharmacological intervention to reduce the levels of both trehalose and glycogen. The lithium cation (Li^+^) inhibits the yeast enzyme phosphoglucomutase (Pgm) by displacing its magnesium cofactor(*25*). Pgm catalyzes the formation of glucose-6-phosphate, a necessary precursor to both trehalose and glycogen, and treatment with LiCl has been shown to reduce levels of trehalose and glycogen accumulated in response to heat (*25*) (Figure 3A). Consistent with this, treatment of cultures with 50 mM lithium chloride prior to and during heat stress reduced accumulation of glycogen by iodine staining (*25*) (Figure 3D). Furthermore, LiCl treatment of cultures growing in steady state at 30°C or 37°C resulted in a disequalization of reaction rates with the reaction now proceeding 50% faster at 37°C, in agreement with the results in cell lysates (Figure 3D). Treatment with an equivalent concentration of NaCl did not affect viscoadaptation, ruling out general effects of salt treatment (Supp Figure 7). Together, these data indicate that accumulation of trehalose and glycogen is the mechanism underlying viscoadaptation.

### Trehalose and glycogen increase viscosity while maintaining solubility of proteins

To investigate the contributions of trehalose and glycogen to viscosity, we measured diffusion in solutions of varying trehalose and glycogen content at 22°C and 40°C using FRAP (Figure 4A). As expected, the presence of either trehalose or glycogen slowed GFP diffusion relative to pure water with a combination of both carbohydrates demonstrating the largest effect. Notably, the diffusion rate was temperature sensitive as expected (faster at 40°C relative to 22°C) for all solutions except 45% trehalose (Figure 4A). Microrheology on a 45% trehalose solution at 22°C and 40°C confirmed that the diffusion rate was insensitive to this temperature difference (Figure 4B). This appears to be unique property of trehalose with likely implications for its role in viscoadaptation.

**Figure 4.**
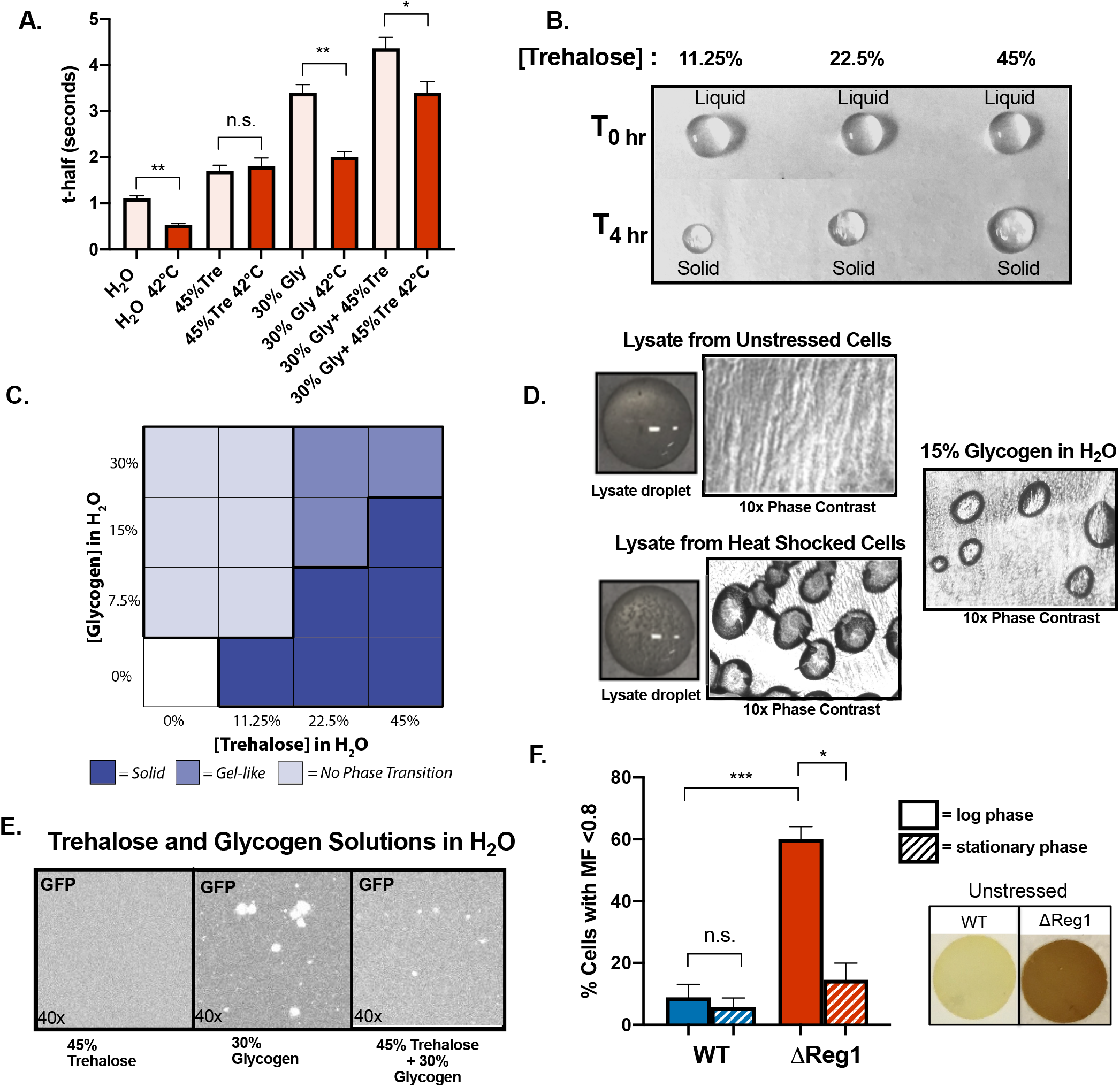
Trehalose and glycogen in combination increase viscosity while maintaining solubility. (**A**) FRAP on purified GFP in aqueous solutions of trehalose and glycogen at 22°C and 40°C. (n≥3, unpaired t-test H_2_O 22°C vs 42°C p=.0023, 30% Gly 22°C vs 30% Gly 42°C p=.0013, Gly/Tre 22°C vs Gly/Tre 42°C p=.0286)(**B**) Aqueous solutions of trehalose at the indicated concentrations either as liquid droplets (top row) or their corresponding solids (bottom row) (**C**) Summary of observations on the phase behavior of trehalose and glycogen droplets of varying concentration at 22°C, phase designation criteria in Supp. Figure 8. (**D**) Droplets of lysate from unstressed or heat shocked cells. Left images are after 10 minutes at room temperature and right images are after 3 hours when lysates have solidified. Far right image is 15% glycogen in H_2_O after droplet solidification. (**E**) Fluorescent images of purified GFP in solutions of trehalose and glycogen. (**F**) WT and ΔReg1 cells were catagorized by GFP mobile fraction (l<0.8 = low, >0.8 = high) in logarithmic growth (unstressed) or stationary phase (nutrient depleted) cultures. (unpaired t-test WT log vs. ΔReg1 log p=.001, ΔReg1 log phase vs ΔReg1 stationary phase p= .0214) (n≥2 experiments, ≥10 cells/experiment). All error bars are s.e.m.

Synthesis of trehalose and glycogen almost always occur together in cells, suggesting that their functions may be complementary(*26*). Cells deficient in glycogen synthesis, but with unimpaired trehalose synthesis, displayed a wide range of viscosities in response to heat, whereas strains deficient in both glycogen and trehalose had a uniform, low viscosity phenotype (Figure 3B). This suggests that glycogen may modify the behavior of trehalose in cells. Because liquid-to-solid phase transitions have been demonstrated for trehalose in certain biological contexts, the hyper-viscous state observed for some cells in the absence of glycogen could be related to phase transition of trehalose(*27*). We examined the effect of glycogen on phase transition of trehalose *in vitro* by observing the behavior over time of liquid droplets containing trehalose, glycogen, or both. Trehalose-only droplets underwent a liquid-to-solid phase transition at a specific concentration, evidenced by the linear relationship between trehalose content of the initial droplet and volume upon solidification (Figure 4B).

Glycogen-only droplets did not undergo phase transition and in general the presence of glycogen in trehalose droplets inhibited their transition to solids, instead promoting a gel-like state (Figure 4C)(Supp. Figure 8). The functional concentration range of trehalose in viscoadaptation is likely narrow since the concentrations that increase viscosity are close to those at which phase transition takes place. Based on these findings, we posit that glycogen widens this functional range by simultaneously increasing viscosity while inhibiting phase transition of trehalose. This model fits with the high variance in viscosities that we observed in heat shocked cells with intact trehalose synthesis but impaired glycogen synthesis.

We previously observed that lysate from heat shocked cells had highly unstable phase properties. Closer examination of droplets of these lysates under phase contrast microscopy revealed an extensive network of ovoid inclusions present selectively in lysate from heat shocked cells. Similar structures were observed in lysates from cells grown at 37°C but they were fewer in number and took longer to appear (on the order of an hour as opposed to within a minute). Glycogen is poorly soluble in water (*22*) and we reasoned that precipitation of glycogen might underlie the formation of these structures. Consistent with this, phase contrast microscopy on aqueous glycogen droplets as they dried over time showed the appearance of qualitatively similar structures (Figure 4D). Notably, the addition of trehalose reduced formation of these inclusions, suggesting that trehalose may improve the solubility of glycogen.

Given the apparent tendency to precipitate from aqueous solution, we asked whether glycogen also affects the solubility of proteins. Looking at purified GFP in a 30% glycogen solution, we observed widespread formation of irregular inclusions and a loss of overall fluorescence intensity relative to water where GFP was entirely diffuse. Trehalose has well-documented roles as a molecular chaperone(*28*). Accordingly, GFP remained entirely diffuse in a 45% trehalose solution. We next tested whether trehalose could modify the formation of GFP inclusions in 30% glycogen. Indeed, we found that GFP in a solution of 30% glycogen plus 45% trehalose had qualitatively reduced inclusion formation and partial restoration of fluorescence intensity compared to 30% glycogen alone (Figure 4E).

Given the propensity of glycogen to cause inclusion formation of GFP *in vitro*, we tested whether high glycogen *in vivo* also affects the solubility of GFP. To do so, we generated a GFP-expressing strain lacking the nutrient-sensing protein Reg1, which results in constitutively high intracellular glycogen and a growth defect which is suppressed by knockout of glycogen synthesis machinery(*29*) (Figure 4F). Loss of GFP solubility in these cells should be reflected by a reduction in the GFP mobile fraction, that is the proportion of GFP that is free to move around the cell, as determined by FRAP. When we performed FRAP on unstressed WT and ΔReg1 cells, we found no consistent difference in t-half values, but the ΔReg1 strain had a substantially higher percentage of cells with low mobile fractions (<0.8) compared to WT (60% vs 8%) (Figure 4F).

While the constitutively high glycogen in ΔReg1 cells is abnormal, certain environmental conditions, such as entry into stationary phase, prompt the adaptive accumulation of similarly high glycogen levels(*30*)(Figure 4F). We therefore asked whether the cellular context of glycogen accumulation (“appropriate” or “inappropriate”) influences its effect on GFP solubility. To investigate this, we compared the percentage of cells with low GFP mobile fractions in stationary phase ΔReg1 cells (“appropriate” context) to unstressed ΔReg1 cells (“inappropriate” context). Stationary phase significantly reduced the percentage of Reg1 cells with low mobile fractions (14.5% vs 60%) whereas there was not a significant difference between stationary phase and the unstressed state for WT cells (5.8% vs 8.9%) (Figure 4F). From these observations we propose that cellular processes that accompany entry into stationary phase, such as production of trehalose, can mitigate the effects of high glycogen on protein solubility. Overall these findings support a model wherein trehalose improves solubility of glycogen and proteins within high glycogen environments, whereas glycogen acts as a viscogen while disfavoring phase transitions driven by trehalose.

### Viscoadaptation occurs in response to glucose limitation

Our results thus far have demonstrated that levels of trehalose and glycogen are closely linked to the diffusibility of proteins in the cell and that this is regulated by temperature. However various forms of nutrient deprivation, including stationary phase and glucose deprivation, induce the accumulation of trehalose and glycogen (*26, 31*). We therefore asked whether glucose deprivation affects diffusion, and thus reaction rates, in cells. Using BirA as a model reaction, we found that glucose limitation reduced the rate of the BirA reaction in cells by about 60% (Figure 5B). Once again, this observation held for a variety of BirA-Protein/Avi-Protein pairs (Supp Figure 9A, 9B, 9C, 9D).

**Figure 5.**
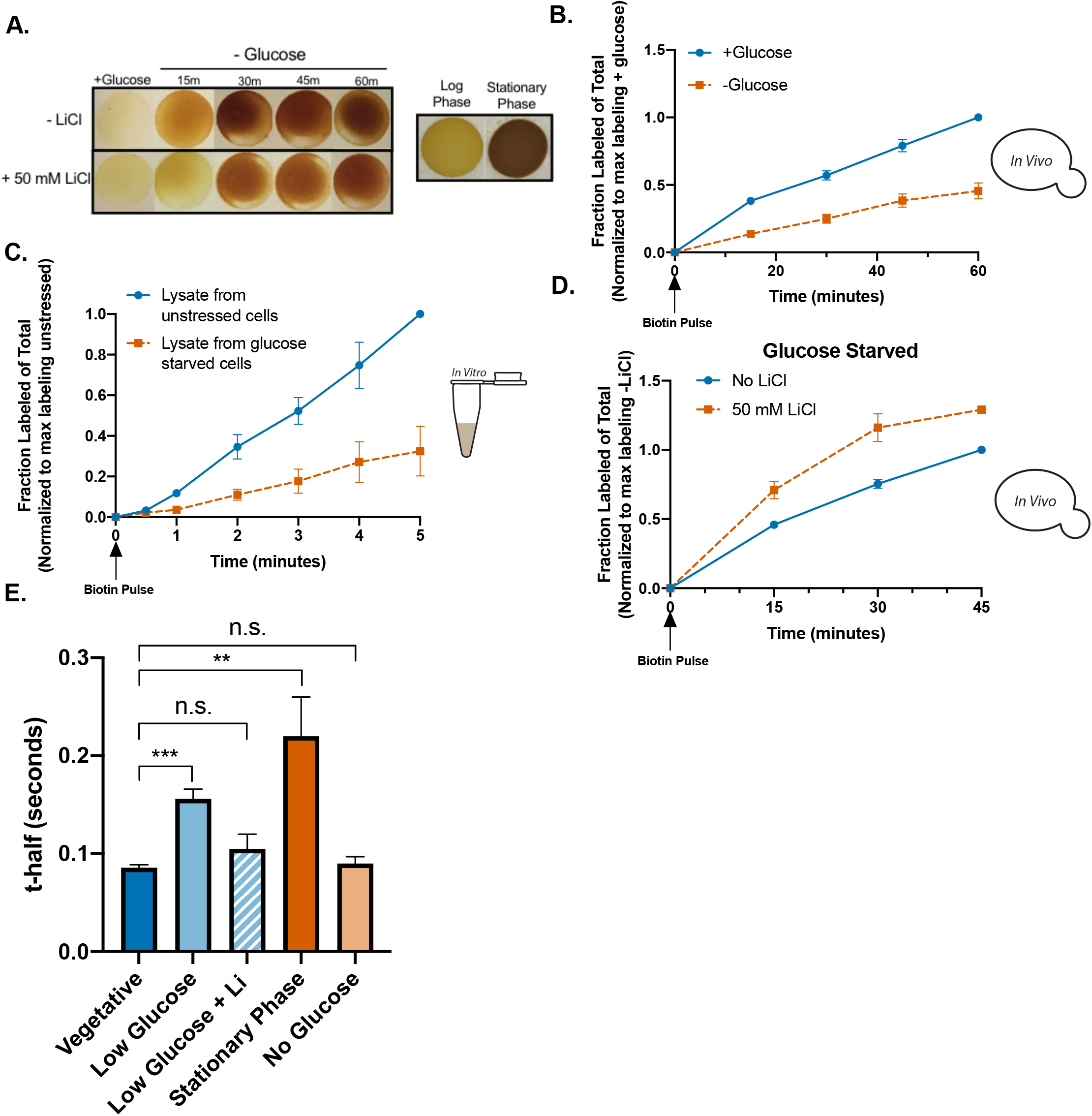
Glucose deprivation and stationary phase cause viscoadaptation. (**A**) Iodine staining on yeast cells glucose starved for the indicated duration with (bottom row) or without (top row) treatment with 50 mM LiCl. Far right panel contains iodine stained yeast spots from logarthmic or stationary phase cultures. (**B**) BirA labeling timecourse *in vivo* with GFP-Avi as “bait”. Reactions in vegetative (+glucose, blue) and starved (-glucose, orange) conditions.(**C**) *In vitro* BirA labeling timecourse in lysates from vegetative (blue) and starved (orange) culturess. (**D**) *In vivo* BirA labeling in glucose starved cells with no drug (blue) or 50 mM LiCl (orange) (**E**) FRAP measurements on cells in the indicated growth conditions (n≥ 3 experiments, ≥10 cells/experiment) (unpaired t-test vegetative vs. low glucose p=.0006, vegetative vs stationary phase p= .0094) All error bars are s.e.m.

The BirA reaction requires ATP so to rule out effects due to changes in ATP, we measured the reaction rate in glucose starved (30 min prior to lysis) and unstarved lysates with an excess of supplemental ATP. The reaction rate was 66% slower in lysate from starved cells relative to unstarved cells, in close agreement with the *in vivo* rate discrepancy (Figure 5C). Furthermore, lithium treatment for one hour prior to starvation reduced glycogen accumulation and increased the reaction rate by 25% relative to untreated cells, suggesting that viscoadaptation is the mechanism of rate regulation (Figure 5C, Figure 5D). We next directly measured diffusion of GFP in glucose limited cells using FRAP. The average t-half for GFP in starved cells was ~0.16s, double that of unstarved cells (Figure 5E). LiCl treatment brought the average t-half back down to 0.1s (Figure 5E). We also measured GFP recovery in stationary phase cells, where trehalose and glycogen are known to be high, and found that the average t-half was 0.24s, 300% the t-half in unstarved cells (Figure 5E).

Of note, cellular responses were different depending on the specific conditions of glucose limitation. Cells transferred to 0% glucose did not respond with viscoadaptation, presumably due to lack of glucose for trehalose and glycogen synthesis. However, when cells were grown in low biotin media, as required for the biotinylation assays, 0% glucose induced viscoadaptation (Supp Fig 10). In low biotin conditions cells poorly metabolize glucose, which results in a constitutive intracellular glucose pool(*32*). We therefore speculate that this residual glucose enables cells in low biotin to initiate viscoadaptation despite 0% glucose. Accordingly, cells in regular media can initiate viscoadaptation in response to 0.1% glucose (Figure 5E). Together these observations reveal that viscoadaptation occurs in response to glucose limited conditions.

### Viscoadaptation occurs in response to low ATP levels and links energy availability to diffusibility of proteins in the cytosol

Both temperature change and nutrient deprivation, two seemingly disparate stressors, may trigger viscoadaptation through a common signal. One commonality between these stresses is rapid loss of intracellular ATP (*33–36*). We therefore asked whether viscoadaptation could be triggered by low ATP. To investigate this, we used pharmacological perturbations that are known to reduce ATP levels. Treating cells with the metabolic poisons sodium azide, sodium arsenite, or a combination of Antimycin A, Oligomycin, and FCCP all increased intracellular viscosity as measured by FRAP (Figure 6A) (*37, 38*). Furthermore, changes in viscosity induced by these treatments were fully or partially prevented by pre-treatment with lithium, suggesting viscoadaptation as the mechanism (Figure 6A).

**Figure 6.**
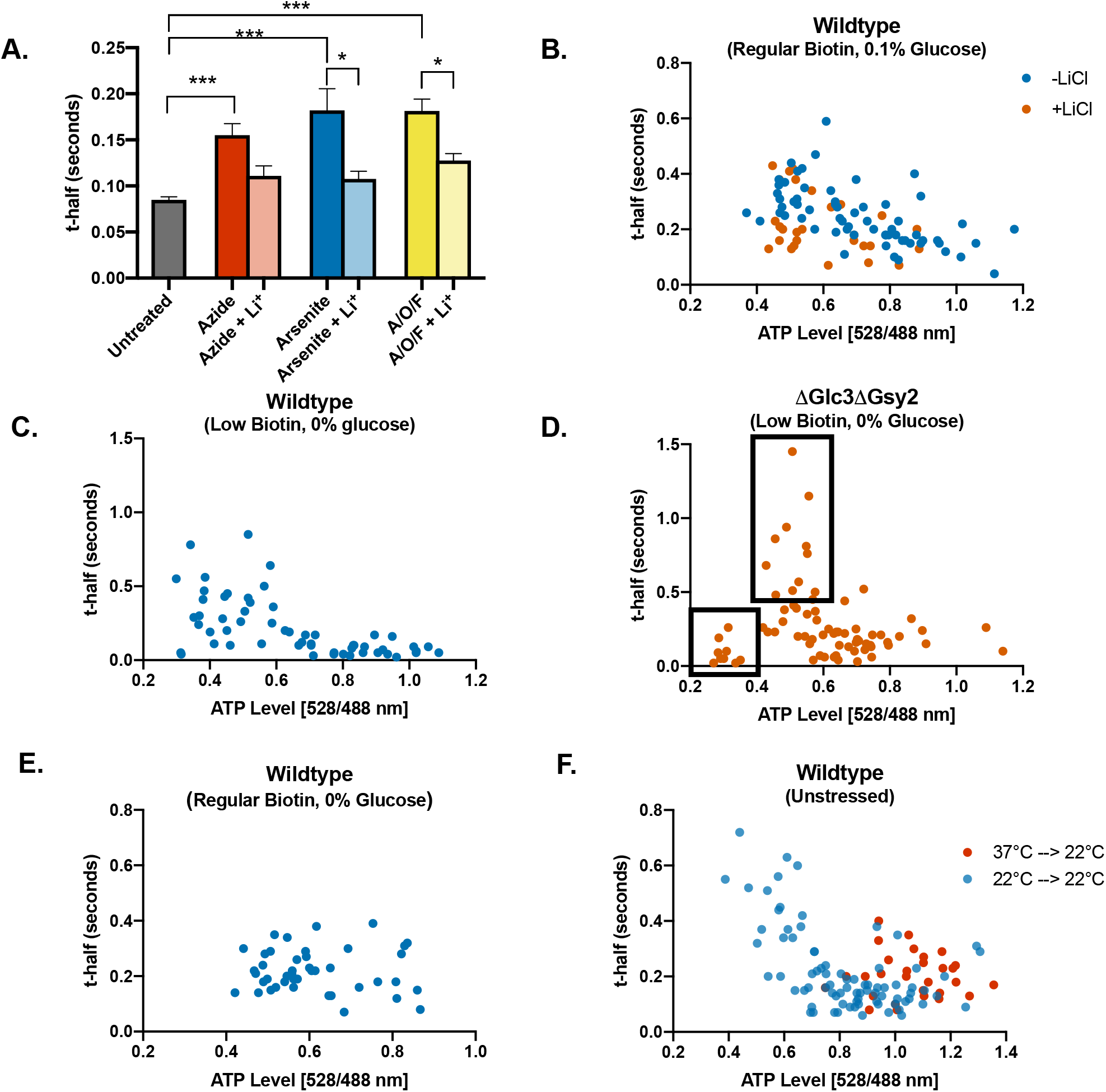
Conditions that cause low ATP induce viscoadaptation. (**A**) FRAP measurements on WT cells treated with NaN_3_, NaAsO_2_, or a combination of oligomycin, antimycin A, and FCCP, with and without the addition of LiCl. (n≥3 experiments, ≥15 cells/experiment) All error bars are s.e.m. (**B**) WT cells expressing a FRET-based ATP nanosensor in 0.1% glucose. ATP levels for individual cells were determined by the ratio of 528:488 fluoresence. Mobility in the same cells was assessed by FRAP on the fluorescent sensor. Cells were treated with or without 50 mM LiCl 1.5 hours prior to glucose limitation (without lithium corr. coefficient = −0.597, p<.0001; with lithium corr. coefficient= −0.4, p=0.0152). (**C**) WT cells in 0% glucose with low biotin (0.5 nM)(corr. coefficient = −0.589, p<.0001). (**D**) t-half values and ATP levels are plotted for the ΔGlc3ΔGsy2 knockout strain in 0% glucose with low biotin (corr. coefficient= −0.132, p=0.135). (**E**) t-half values and ATP levels are plotted in WT cells in 0% glucose and normal biotin levels (corr. coefficient = −0.047, p=0.38). (**F**) Cells growing in unstressed conditions (blue points) are sampled based on natural variance in ATP levels and plotted with their respective t-half values (corr. coefficient = −0.61, p<.0001). Red points correspond to cells growing at 37°C which were downshifted to 22°C for measurement.

To examine the relationship between viscosity and ATP at the single cell level, we made use of a genetically encoded FRET-based ATP sensor for which the ratio of 528:488 nm emission is indicative of the cellular ATP level (Supp. Figure 11A, 11B, 11C)(*33*). Cells with high ATP have a higher 528:488 ratio than those with lower ATP(*33*). By performing FRAP measurements on the fluorescent ATP sensor, we were able to measure ATP and diffusion in the same cells (Figure 6A) (*39*)(*33*). Plotting t-half values against ATP levels (528/488 nm revealed a negative correlation between ATP content and recovery time (Figure 6B). This held true for cells in 0% glucose with low biotin or in 0.1% glucose with regular biotin (r= −0.59 and r= −0.6, respectively), but was not true for cells in 0% glucose with regular biotin, when viscoadaptation does not occur (r= −0.0474). In addition, we found that treatment with lithium prior to the switch to 0.1% glucose reduced the strength of this correlation (r= −0.4). These observations indicate that ATP levels and intracellular diffusion are strongly correlated in conditions where viscoadaptation takes place.

If low ATP triggers viscoadaptation, then strains with impaired viscoadaptation should have an altered relationship between ATP and viscosity. To test this, we performed FRAP on the ATP sensor in glycogen-deficient cells. We observed the emergence of two new populations: cells with low ATP and high mobility (non-responders), and cells with low ATP and extremely low mobility (hyper-responders). These populations mirror the phenotypic variability in viscosity observed for glycogen deficient strains (Figure 3C). Notably, hyper-responders had universally higher ATP than non-responders, suggesting a potential role for viscoadaptation in energy conservation. Prior work has demonstrated that ATP can act as a hydrotrope, helping to keep proteins soluble in cells (*40*). This in turn could affect protein mobility. However, non-responder cells with low ATP and fast diffusion demonstrate that low ATP is not sufficient to reduce mobility.

If viscoadaptation occurs as a direct response to low ATP, we should find that stochastic variation in ATP levels in the absence of any stress is sufficient to confer correlation with intracellular diffusion. While the majority of cells in an unstressed culture have high ATP, we sampled cells with qualitatively different ATP levels to capture the range of natural variation. By performing FRAP measurements on these cells, we again found a strong negative correlation between viscosity and ATP content, that is cells with low ATP had slower GFP recovery times (r= −0.6, p<.0001) (Figure 6F). We next asked whether the relationship between the conformation of the ATP sensor and the viscosity of the cell reflects differences in ATP levels rather than direct stabilization of the “open” (low ATP) conformation under high viscosity. To differentiate between these possibilities, we examined the ATP sensor in conditions predicted to have both high ATP and high viscosity. To do this, we shifted cells from steady state growth at 37°C, where higher viscosity compensates for the higher growth temperature, to a measurement temperature of 22°C, thereby creating a temporary mismatch between the cellular adaptation state and the environment. Measuring cells within the first 15 minutes of switching temperatures resulted in cells with high ATP but lower mobility (orange points) (Figure 6F). We therefore conclude that the sensor maintains function at these lower viscosities. Collectively these findings support the model that low ATP triggers viscoadaptation.

## Discussion

Taken together, our findings reveal a strategy, viscoadaptation, by which cells can broadly modulate intracellular diffusion in response to changes in temperature or under low ATP conditions (i.e starvation, inhibition of mitochondrial function, and stochastic drops in ATP). While a single cell cannot control many aspects of its environment, including temperature, it uses the production of trehalose and glycogen as an orthogonal strategy to control intracellular viscosity, thereby enabling the homeostatic regulation of viscosity-sensitive processes across multiple growth temperatures. In this way, viscoadaptation defines a new role for trehalose and glycogen, two molecules long considered to be primarily “storage” carbohydrates, and reveals viscosity to be a finely tunable cellular property.

Changes in viscosity have the potential to impact diverse cellular processes, including the rates of diffusion-controlled reactions, the temporal regulation of signaling cascades, the accuracy of kinetic proofreading mechanisms, and the assembly or disassembly of large multimeric complexes. Therefore, the ability to regulate viscosity in a manner independent of environment may have evolved as a broad adaptive strategy. For instance, slowing cellular functions on a large scale might allow energy conservation when nutrients are scarce. Trehalose has been implicated in a similar role during dormancy in plant spores where metabolic activity drops to nearly zero, yet spores retain viability(*27*). Under conditions of low ATP, viscoadaptation may represent a more tunable version of this response whereby cells can reduce energy expenditure while maintaining most cellular functions.

Additional mechanisms for regulating the mobility of biomolecules in cells have been described by others, including crowding-based mechanisms, the assembly of stress-induced condensates, and pH-induced protein polymerization. (*41–44*). Some conditions that induce viscoadaptation may also induce these responses. However, viscoadaptation is phenomenologically and mechanistically distinct. Of these responses only viscoadaptation affects mobility on the size scale of individual proteins or implicates trehalose and glycogen in the control of protein diffusion. In addition, viscoadaptation is unique in serving as both an acute response to sudden changes in the environment and a homeostatic mechanism. Moving forward, it will be interesting to see if and how other cellular stress responses interact with the viscoadaptive response.

## Acknowledgements

We thank J. Ferrell, P. Geiduschek, P. Harbury, D. Herschlag, K.C. Huang and R. Das for invaluable guidance in the preparation of this manuscript. All members of the Brandman lab provided helpful discussions and continuous support throughout this project.

## Materials/Methods

### Yeast strains and growth conditions

All yeast strains were in the BY4741 S288C background. Strains were grown at the indicated temperatures in synthetic complete or dropout media with 2% glucose according to the recipe provided by Hanscho et al. (*45*). For biotinylation experiments yeast were grown in low biotin SD complete media with 2% glucose (Sunrise Science-biotin YNB supplemented with 0.5 nM biotin).

Deletion strains were constructed via transformation with PCR products containing antibiotic resistance cassettes (*NATMX6, HYGMX6*, or *KANMX6*). PCR products contained 40bp of homology to the 5’ and 3’ genomic regions immediately adjacent to the gene to be deleted. Cytosolic BirA was integrated at the LEU2 locus driven by a PGK1 promoter. BirA fusion proteins contain a 120 residue, intrinsically disordered linker. Transformants were verified by genomic PCR.

Plasmids used in this study were cloned by the Gibson Assembly method using NEBuilder HiFi DNA Assembly Master Mix (New England Biolabs).

### Biotin Labeling Time Courses

For *in vivo* biotinylation reactions, cells expressing BirA under the PGK1 promoter plus an Avi -tagged substrate under a variable promoter were grown in low biotin media (0.5 nM biotin) to OD 0.4 – 0.6. The genomically integrated BirA strain and the BirA tagging plasmid were a gift from Calvin Jan at Calico. Cultures were subjected to various treatments as described. Addition of biotin to a final concentration of 1 μM initiated the reaction. All samples were corrected for background labeling (prior to the biotin pulse) by comparison to a no biotin control sample. Cells were collected by spinning down 1 ml of OD 0.4-0.6 culture, removing the media and resuspending in 12-15 ul 4x NuPage LDS Sample Buffer with 5% β-mercaptoethanol. Samples were immediately boiled at 95°C for five minutes to ensure denaturation.

For *in vitro* biotinylation assays, lysates from cells expressing the desired Avi-tagged substrates were prepared as described above. *In vitro* labeling experiments were conducted using the Avidity BirA biotin-protein ligase standard reaction kit (Avidity BirA500). Samples were collected at the indicated time points and the reaction stopped by resuspension in 6 ul of boiling 4x NuPage LDS Sample Buffer with 5% β-mercaptoethanol. Samples were then immediately boiled as above.

### Gel Shift Assay

Samples for quantification of biotinylation were allowed to cool for at least 10 minutes after boiling. 6 ul of boiled sample was combined with 1.5 ul streptavidin (Thermo Fischer #S-888 10 ug/uL), briefly vortexed and loaded into a invitrogen SDS-PAGE 1.5 mm 4-12% Bis-Tris gel. Gels were run for 2 hours at 110 V at 4°C. Transfer was performed onto nitrocellulose membrane using a semi-dry transfer apparatus (Bio-Rad Turbo Blot). Membranes were blocked for 30 minutes in 5% milk in TBST and then incubated with mouse anti-HA primary antibody (antiHA-12C5 Roche #11583816001) in 5% milk in TBST with gentle shaking overnight at 4°C. After primary antibody incubation, membranes were washed three times in TBST for a total of 15 minutes and incubated at room temperature with Anti-mouse 800 secondary (1:5000, Licor) in 5% milk in TBST for 1 hour. Blots were imaged on a Licor gel imager and band intensities were quantified using ImageStudioLite. All ratiometric analyses are the intensity of the top band divided by the intensity of the bottom band plus the top band in a given gel lane.

### Cell Lysate Preparation

Cell lysates were created by cryogenic lysis of log phase “flash frozen” yeast (separated from growth media by filtration and immediately suspended in liquid nitrogen). Frozen lysates were ground into a fine powder using a Spex 6750 Freezer Mill at rate 10 for 1 minute. The powder was allowed to thaw with gentle shaking at 4°C, followed by centrifugation at 10,000 × g for 10 minutes at 4°C. Supernatant was removed and subjected to a second spin at 16,000 × g for 10 minutes at 4°C. Supernatant was once again removed and used directly in lysate experiments or stored at −80°C for later use.

### Microrheology

Microrheology experiments were performed on a Ziess LSM 880 confocal microscope using an oil immersion 40x objective. Lysates were mixed with 0.5 um fluorescent beads (488 nm/512 nm). Chamber slides were prepared using double sided tape to delineate the chamber width (1 cm) and depth (100 um). After loading lysate with beads (~20 ul) into the chamber, open ends were sealed with tape to prevent evaporation. Focal plane was chosen to be halfway between the slide and coverslip (50 um from the slide) and X and Y positions were centered in the chamber to avoid boundary effects. 500 images were collected over a total of 30 seconds at 5x zoom using a laser power of 6%.

Particle tracking was done using Particle Tracker 2D/3D in the imagej Mosaic plug-in. Particle detection settings were as follows: Radius 3, Cutoff .001, Per/Abs 0.15. Particle linking parameters were as follows: Link Range 2, Displacement 10, Dynamics: Brownian. Trajectories ≤ 10 frames were eliminated from the analysis. Diffusion coefficients extracted from multiple trajectories in the same frame were averaged.

### Heat Shock

For small cultures (<10 ml) heat shock experiments were performed by pelleting cultures, removing supernatant, and resuspending in media heated to the indicated temperature. Cultures were then placed in an incubator of the appropriate temperature for the duration of the heat shock treatment.

For heat shock of large cultures (≥1 L) media was separated from cells by vacuum filtration onto nitrocellulose membrane. Cells were quickly scraped from the nitrocellulose and resuspended in preheated media. Flasks were then shaken for 30 minutes in pre-warmed incubators.

For heat shock FRAP experiments, room temperature cells were placed on slides and entered into a microscope chamber pre-warmed to the indicated temperature.

### Fluorescence Recovery After Photobleaching (FRAP)

FRAP experiments were carried out on a Ziess LSM 880 confocal microscope using a 40x oil immersion objective. For *in vivo* measurements, 100 images were taken at 11x zoom where bleaching occurred after the second image. Images were separated by 52 milliseconds. Per cell, one bleach zone, one control zone within the cell, and one background zone outside the cell were specified (all circular). Bleach and control zones were generally chosen such that the circle touched one edge of the cell. Regions were selected to be homogenous in their fluorescence intensity. Except for experiments where the mobile fraction was being compared between conditions, any measurement with a mobile fraction below 0.8 was not included in analyses.

For *in vitro* measurements, we used purified GFP (Chromtek, eGFP-250) at a concentration of .17 mg/mL (2 μL GFP into 10 μL lysate). For lysate and solution measurements, between 250 and 1000 images were taken at 1x zoom, depending on the speed of recovery. Bleaching occurred after the second image using 100% laser power in the 405 nm, 456 nm, and 488 nm channels. Images were separated by 120 milliseconds. Bleach zones, control zones, and background zones were circular.

FRAP images were analyzed using the imageJ bioformats package to extract intensity values for each region at each time point and easyFRAP, an open-source(*46*) online FRAP analysis program, was used to calculate the mobile fraction, t-half, and R-squared values for each cell. Gap ratio and bleaching depth were used to determine the quality of the measurement. Images with a gap ratio below 0.6 or a bleaching depth below 0.3 were not included in the analyses.

### Drug Treatments

Cycloheximide (Calbiochem #239764) was used at a final concentration of 50 ug/ml. Sodium azide (Sigma #2002) and sodium arsenite (Sigma #7400) were used at final concentrations of 0.15 mM and 2 mM. Antimycin A (Sigma #A8674), oligomycin (Sigma #75351), and FCCP (Cayman Chemical #15218) were used at 10 μM, 0.1 μM, and 50 nM, respectively. Cycloheximide was added 10 minutes prior to the start of any experiment; all other drugs were added 30 minutes prior to the start.

### Iodine Staining

Iodine staining experiments (see Figure 3C) were adapted from the method described by *Ruiz et al* (*29*). Briefly, experiments were performed by pipetting 1 ml of log phase (OD 0.4 – OD 0.6) culture onto a square of nitrocellulose membrane resting on top of a vacuum flask. A circular plastic stencil was used to delineate the area of each spot. The nitrocellulose membrane was then placed in a chamber containing iodine crystals, but not touching the iodine, and allowed to stain for 10 minutes. Qualitative differences in glycogen content were determined by the color of each spot after staining, with darker spots corresponding to cells with more glycogen.

### *In Vitro* Glycogen and Trehalose Phase Experiments

Trehalose and glycogen solutions of the indicated concentrations were made in water using glycogen from oyster (Sigma #G8751) and trehalose dihydrate (EMD Millipore #625625). 40 uL droplets were made on parafilm at room temperature and left 8 hours, by which point trehalose-only droplets had solidified. Phase comparisons were done qualitatively by observing the response of the droplet to application of force with a pipette tip; droplets that dried out without retaining their 3-dimensional structure were considered not to have undergone phase transition. Droplets that distorted upon application of mechanical force without fracturing were categorized as gel-like, and droplets that fractured with application of mechanical force were categorized as solids (see Supp. Figure 8 for examples).

### Glucose Starvation

For glucose starvation experiments yeast were spun down in a tabletop centrifuge at 14,000 × g for 1 minute. Media was discarded and the cell pellet was washed once with glucose-free SD media, before being resuspended in glucose-free or low glucose (0.1%) SD media. Cultures were placed in a spinning wheel at 30°C for the duration of starvation.

### ATP Measurements

ATP levels in single cells were measured using a FRET based nanosensor expressed from a cen/ars plasmid, pDR-GW AT1.03YEMK, which was a gift from Wolf Frommer. (Addgene plasmid # 28004 ; http://n2t.net/addgene:28004 ; RRID:Addgene_28004). Single-cell FRET measurements were conducted on a Ziess LSM880 confocal microscope. Excitation was provided at 405 nm and emissions were detected at 485 (detector settings 460 nm-510 nm) and 535 nm (detector settings 520 nm-560 nm). Measurements were calculated as a ratio of 535:485 signal intensity. The detector gain was set at 750 for the 535 nm channel and 650 for the 485 nm channels, such that the 535 nm:485 nm ratio was roughly 1:1 in unperturbed cells. Measurements were taken following some perturbation to this initial state.

---

**Supplementary Figure 1.**
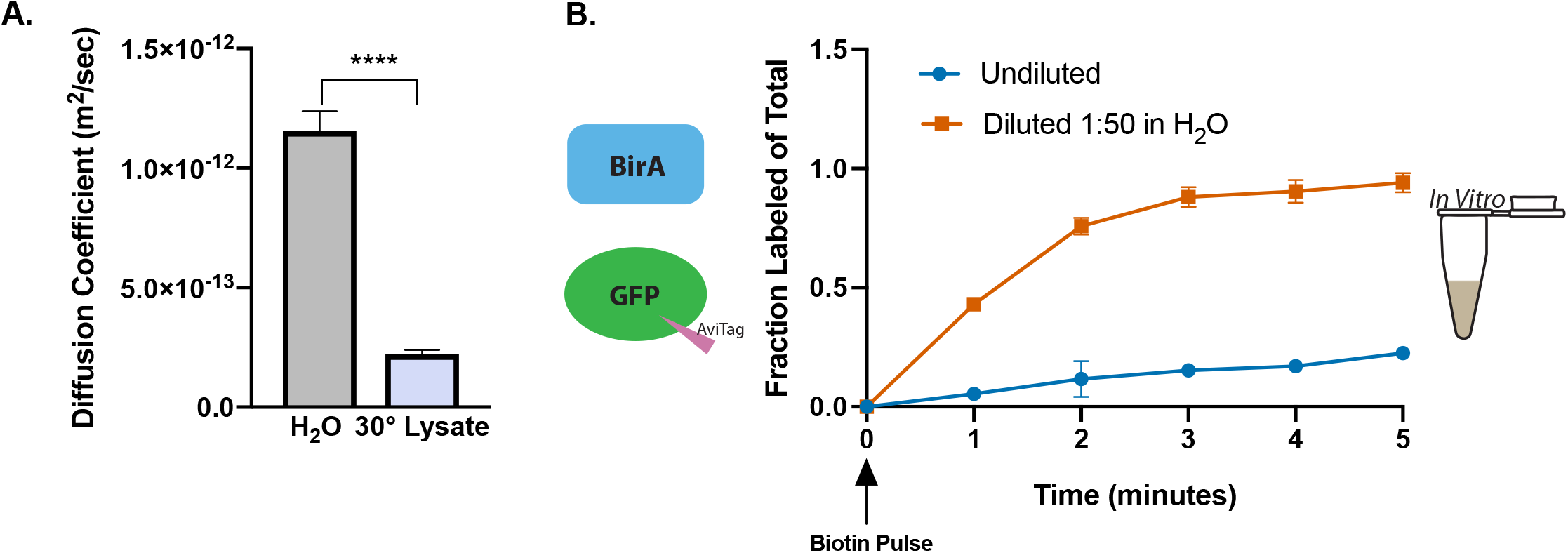
Dilution of cell lysate with water decreases its viscosity and increases the rate of the BirA reaction. (**A**) Fluorescent bead microrheology to measure diffusion coefficients in diluted (1:50 in H_2_O) and undiluted cell lysates. (unpaired t-test p<.0001) (**B**) Rate of *in vitro* GFP-Avi biotin labeling by BirA in diluted (1:50 in H_2_O) (■) and undiluted (●) cell lysates. All error bars are s.e.m.

**Supplementary Figure 2.**
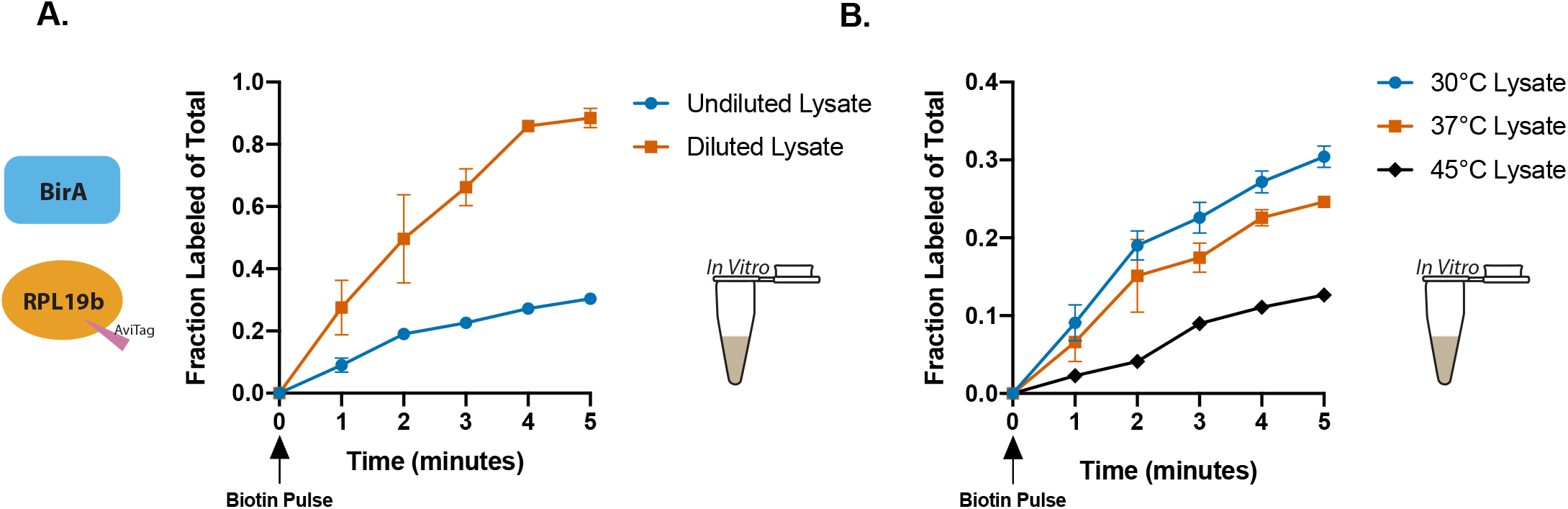
Labeling of RPL19b-Avi by cytosolic BirA is fastest in highly diluted lysate and slowest in undiluted lysate from heat shocked cells. (**A**) Rate of RPL19b-Avi biotin labeling by BirA in undiluted (●) and diluted (1:50 in H_2_O) (■) lysates. Biotin added at 0 minutes. (n=4) (**B**) Rate of RPL19b-Avi labeling by BirA in lysates produced from cells growing at 30°C (●), 37°C (■), or heat shocked for 30 minutes at 45°C (◆). All reactions preformed at room temperature. Biotin added at 0 minutes. (n=4) All error bars are s.e.m.

**Supplementary Figure 3.**
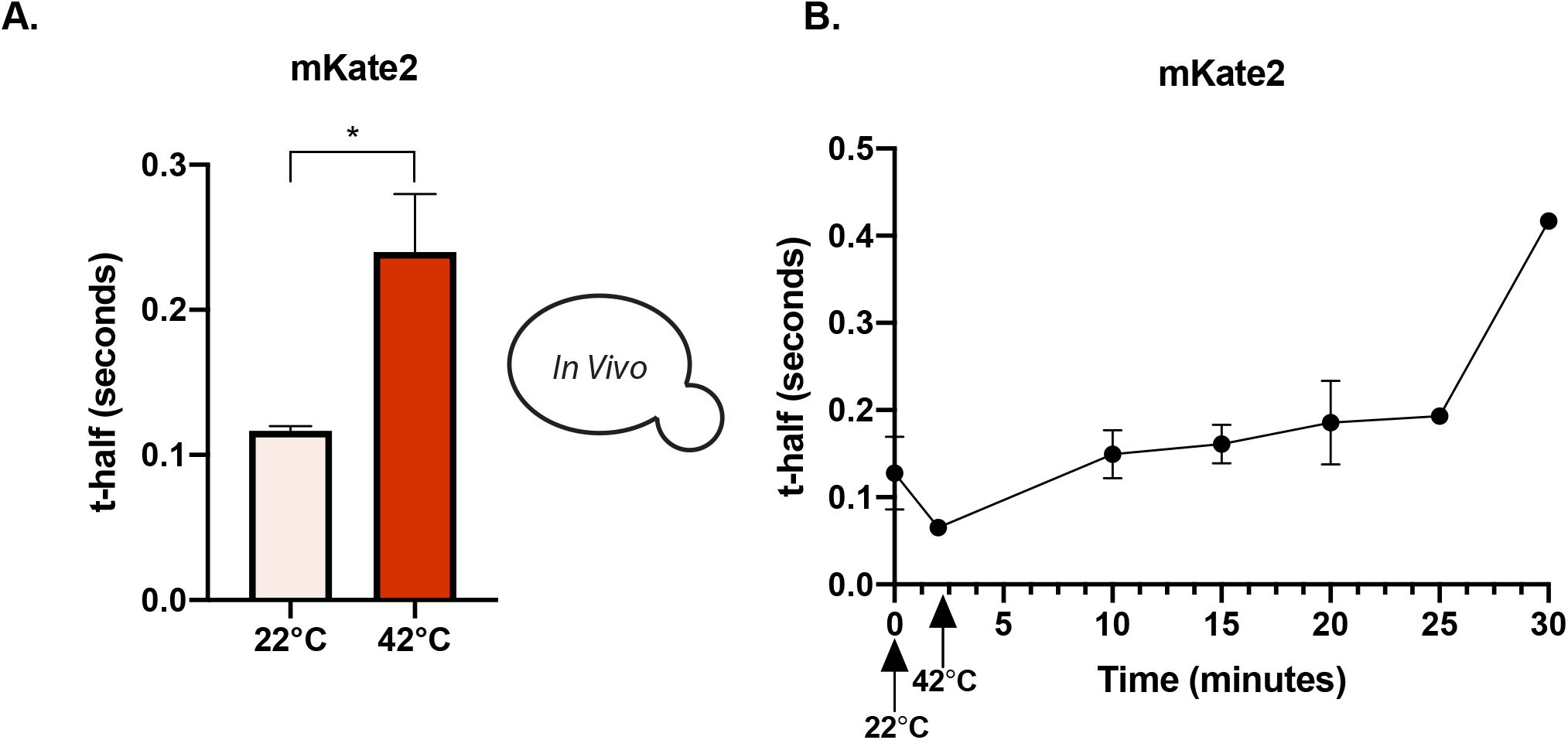
Increased temperature slows diffusion of mKate2 in cells. (**A**) FRAP measurements on cells expressing mKate2 were performed 20 minutes after cells were moved from room temperature (22°C) to a confocal microscope chamber of the indicated temperature. (n=3 cultures/temp, <10 cells/ culture) (unpaired t-test, p=.0271) (**B**) Timecourse of FRAP measurments on cells expressing mKate2. Time 0 corresponds to measurments taken prior to temperature shift (22°C to 42°C). All subsequent time points are after the temperature shift. (n=3 experiments, ≥ 3 cells/time point/experiment). All error bars are s.e.m.

**Supplementary Figure 4.**
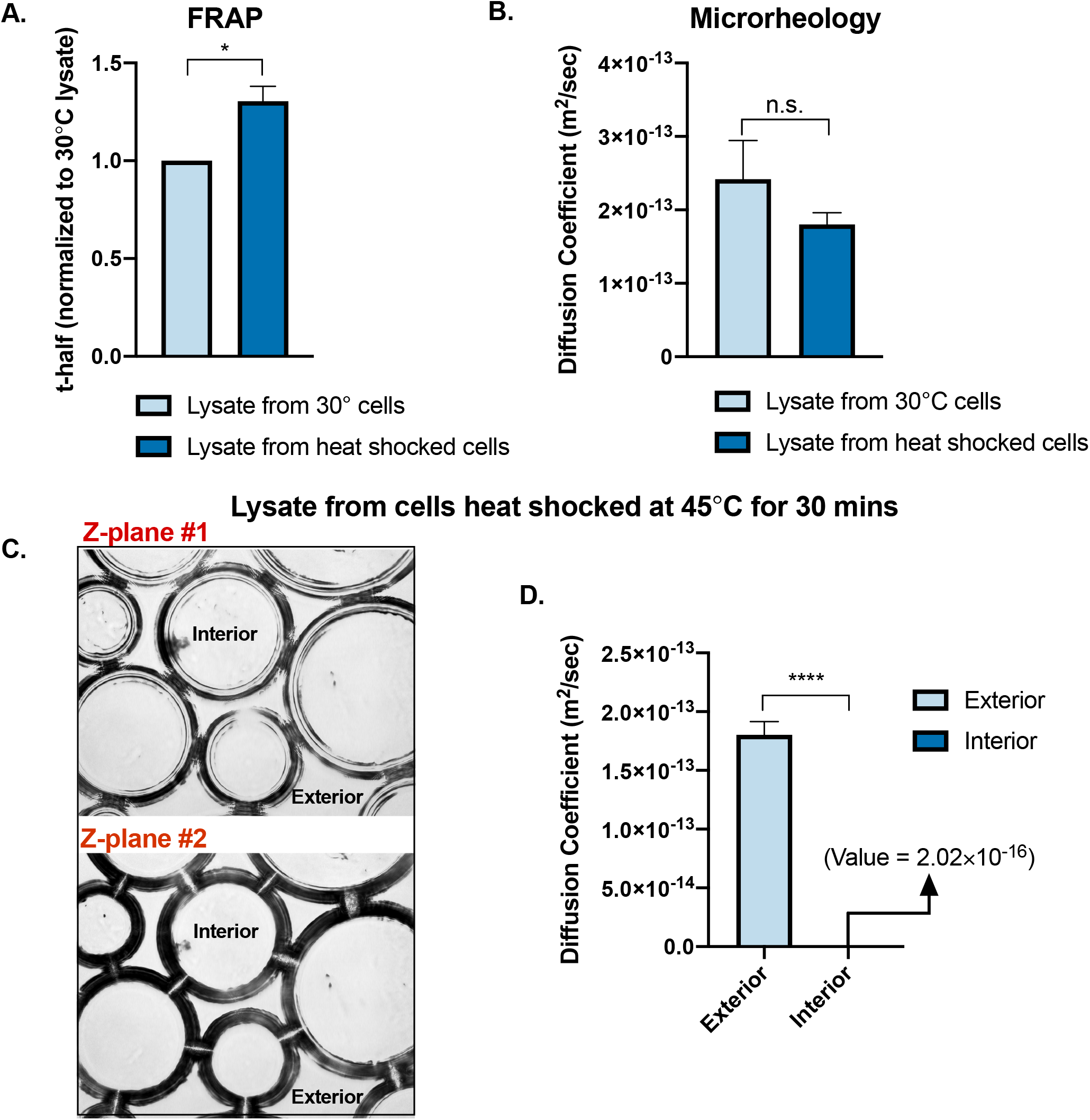
Lysates from heat shocked cells rapidly separate into mobile and immobile regions when water is allowed to evaporate from the lysate. (**A**) FRAP on purified GFP added to lysate from cells grown at 30°C (left) or cells heat shocked at 45°C for 30 minutes (right). Measurments were obtained from mobile regions of the lysates. (n=3 lysates/condition)(paired t-test, p=.02) (**B**) Microrheology of fluorescent beads in the indicated lysates. Measurments were obtained from mobile regions of the lysates. (n≥2 lysates/condition) (unpaired t-test p=.17)(**C**) 10x brightfield microscopy on lysate from cells heat shocked at 45°C for 30 minutes. Top and bottom images are two different z-planes of the same region. (**D**) Microrheology measurements on fluorescent beads in regions exterior to the inclusions and interior to the inclusions. (paired t-test p<.0001) All error bars are s.e.m.

**Supplementary Figure 5.**
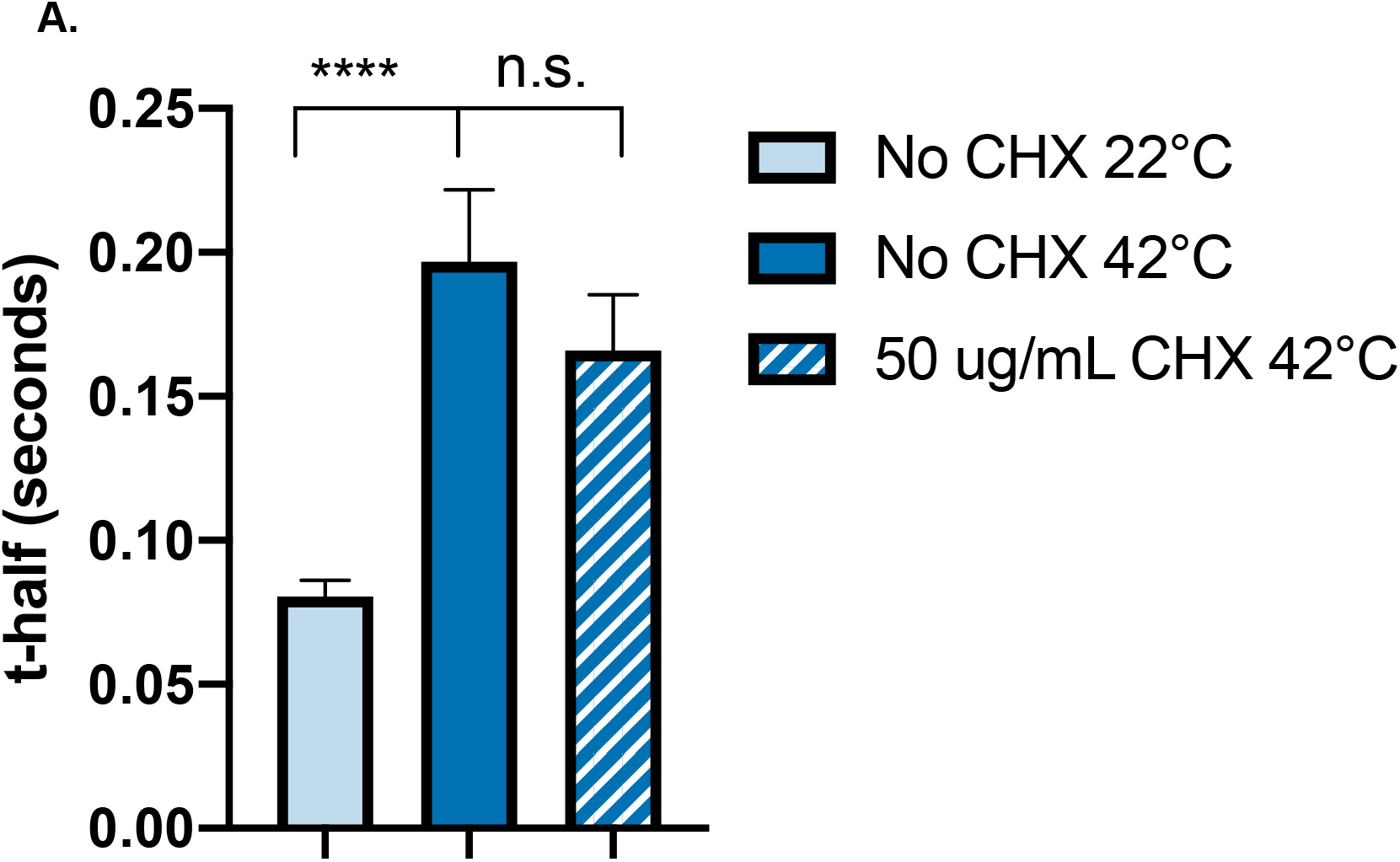
Viscoadaptation in response to acute heat shock does not require protein synthesis. (**A**) Cells were treated with 50 ug/ml cycloheximide (CHX) or no drug for 10 minutes prior to and during heat shock at 42°C. FRAP measurements were performed beginning at 20 minutes after temperature shift. (unpaired t-test on “No CHX 22°C” vs “No CHX 42°C”, p= <0.0001)(unpaired t-test on “No CHX 42°C” vs. “50 ug/mL CHX 42°C”, p=0.169) Error bars are s.e.m.

**Supplementary Figure 6.**
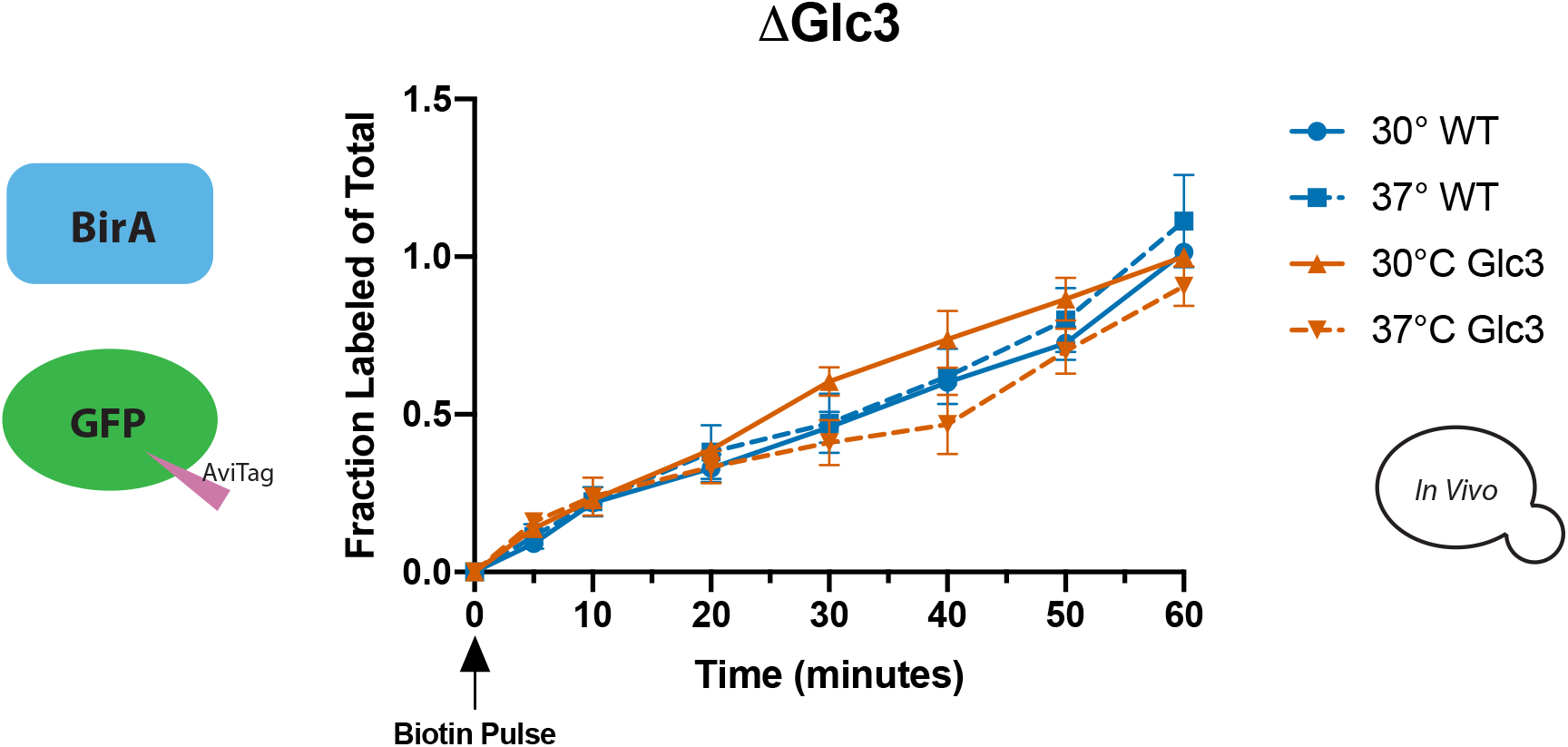
Impairing glycogen synthesis does not change the labeling rate of GFP-Avi by BirA at 30°C or 37°C. (**A**) Wildtype cells (blue) and cells lacking the glycogen branching enzyme Glc3 (orange) were grown at 30°C (solid lines) and 37°C (dashed lines). Biotin labeling was initiated at time 0. (n=4 for WT, n=2 for ΔGlc3) Error bars are s.e.m.

**Supplementary Figure 7.**
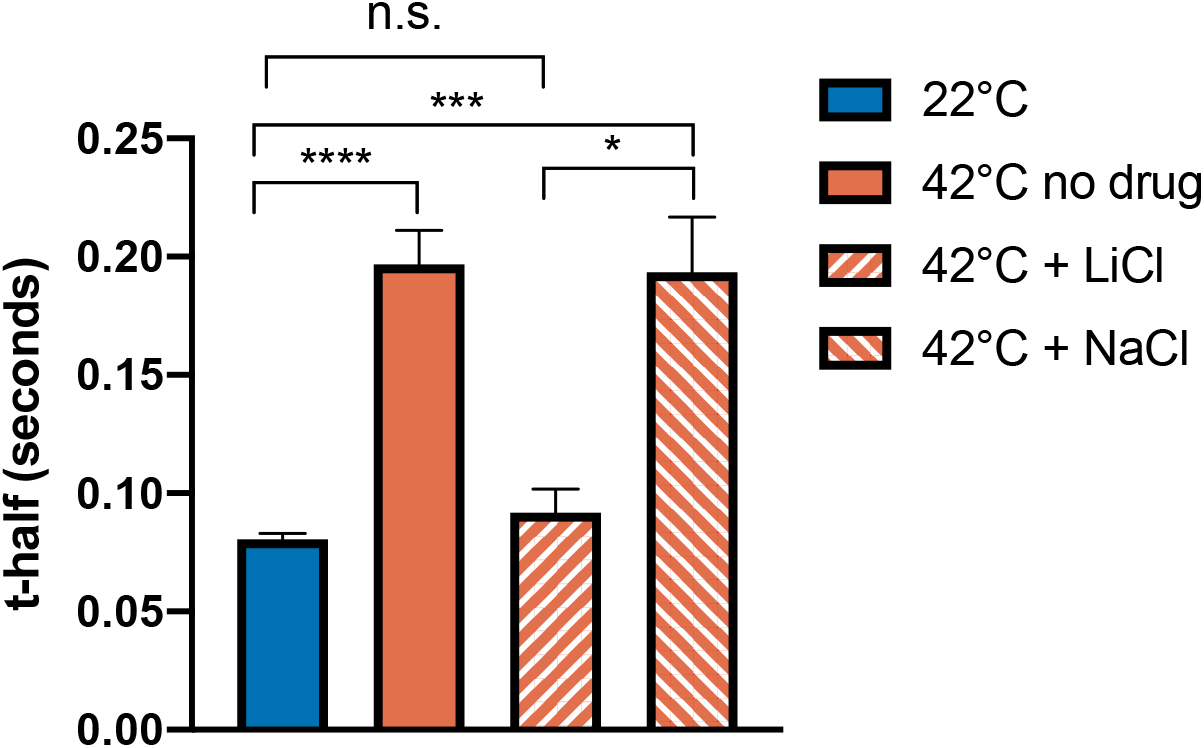
Treatment with 50 mM NaCl does not impair viscoadaptation in response to heat shock at 42°C. Cultures were treated with no drug, 50 mM LiCl, or 50 mM NaCl for one hour prior to temperature shift. FRAP measurements were collected beginning at 20 minutes after temperature increase. Error bars are s.e.m. (n≥ 2 experiments, >10 cells/experiment)(unpaired t-test 22°C vs 42°C no drug p<.0001, 22°C vs 42°C + NaCl p=.0003, 42 + LiCl vs 42 + NaCl p=.018)

**Supplementary Figure 8.**
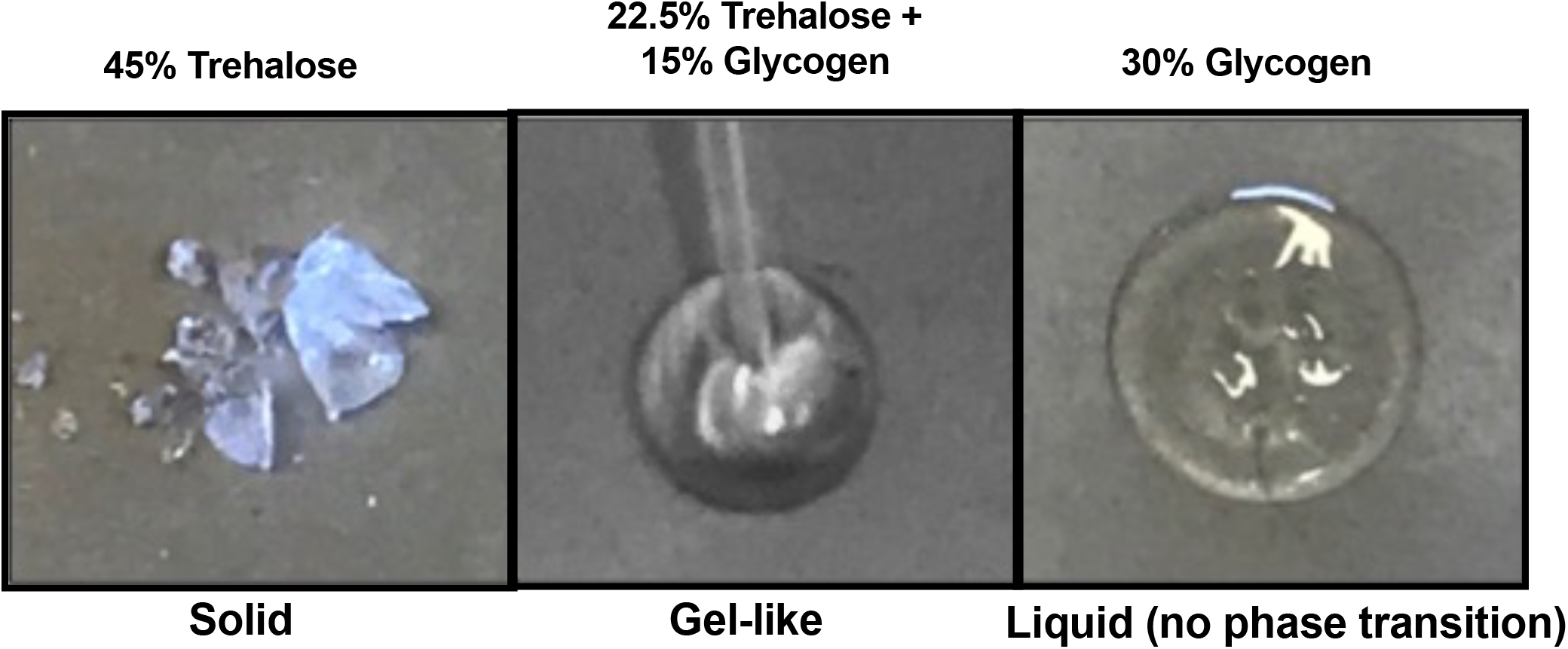
Examples of trehalose and glycogen droplets in different phases. Aqueous droplets of trehalose and glycogen were monitored over time for phase changes. Phase properties of all droplets were assessed once trehalose-only droplets had formed solids. Droplets that fractured in response to mechanical force (left panel) were characterized as “solids”. Droplets that were characterized by distortion upon mechanical force without fracture (middle panel) were considered “gel-like”, and droplets that did not undergo any phase transition were characterized as “liquids” (right panel). Droplets that remained liquids showed continuous evaporation of water leaving behind a solid crust that did not retain the 3-dimensional shape of the initial droplet.

**Supplementary Figure 9.**
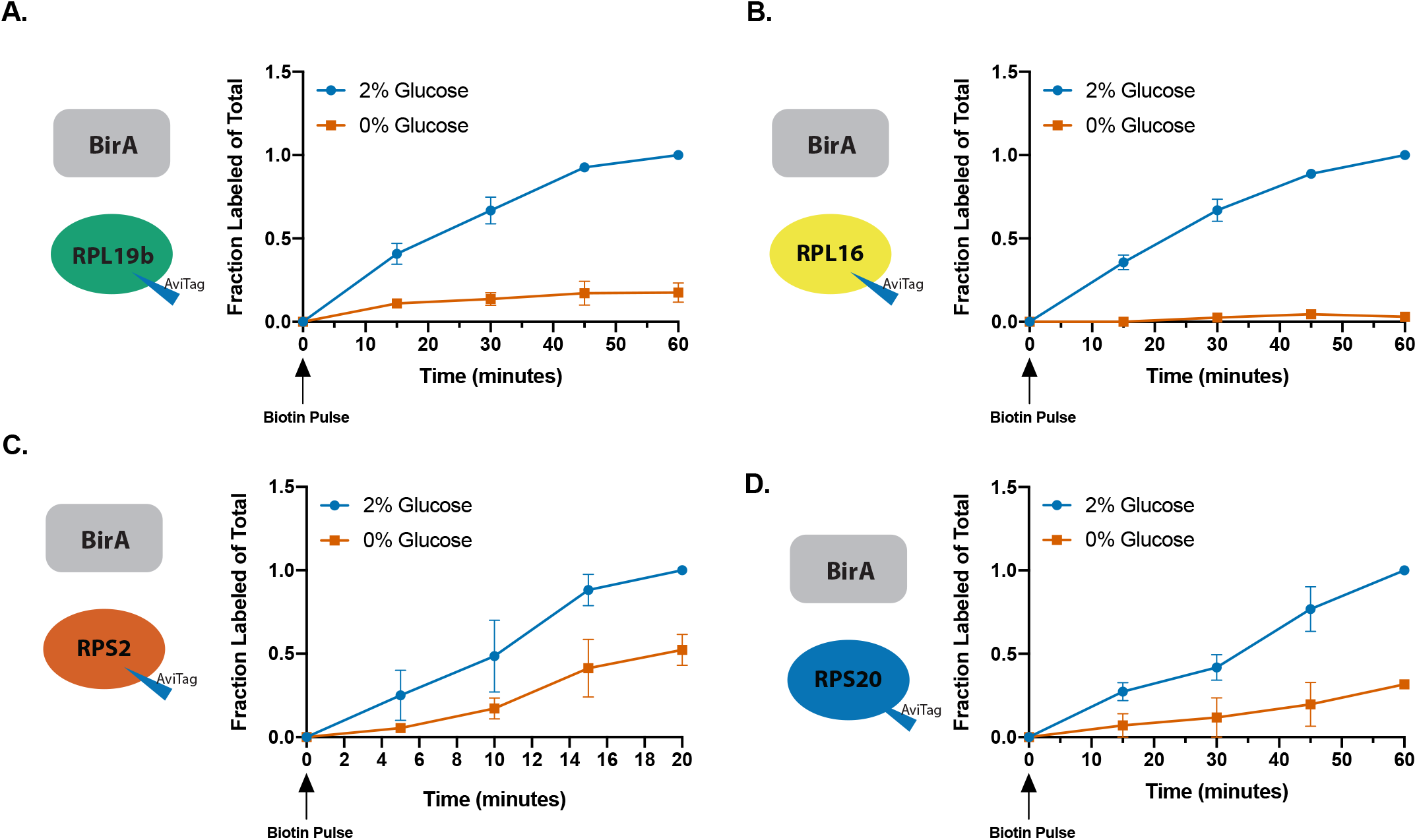
Glucose limitation slows down BirA labeling *in vivo* for multiple Avi-taggged ribosomal proteins. (**A**) Labeling of RPL19b-Avi by cytosolic BirA in 2% glucose (●blue) and 0% glucose (■orange) conditions. Cells are grown in low biotin media (0.5 nM) to prevent background labeling. (**B**) Labeling of RPL16-Avi by cytosolic BirA in 2% glucose (●blue) and 0% glucose (■orange) conditions. (**C**) Labeling of RPS2-Avi by cytosolic BirA in 2% glucose (●blue) and 0% glucose (■orange) conditions. Timecourse shortened due to faster labeling of this Avi substrate. (**D**) Labeling of RPS20-Avi by cytosolic BirA in 2% glucose (●blue) and 0% glucose (■orange) conditions. All error bars are s.e.m.

**Supplementary Figure 10.**
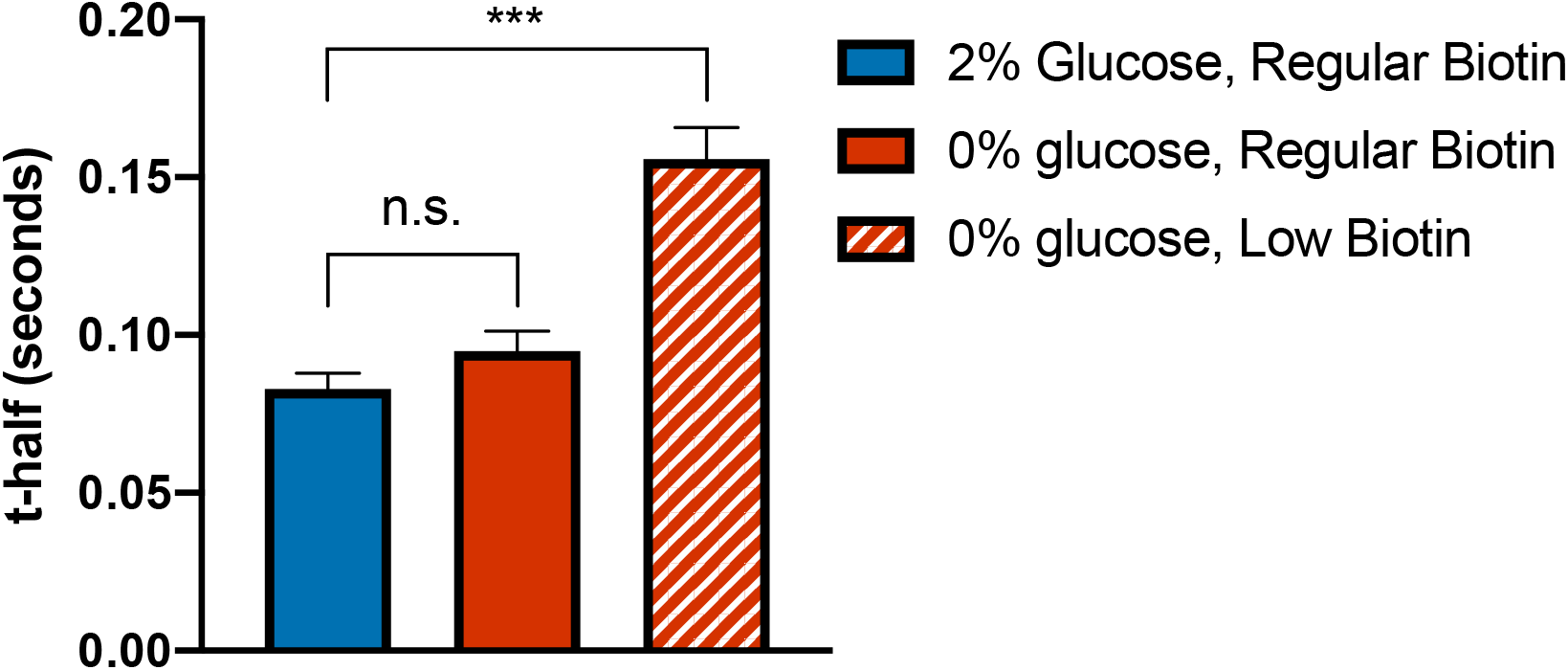
Cells undergo viscoadaptation in response to 0% glucose only when grown in low biotin media prior to starvation. (**A**) Cells were grown in 2% glucose with normal biotin (2 ug/L) or low biotin (.125 ug/L) and transferred to 0% glucose as indicated. FRAP measurements began 30 minutes after removal of glucose. (n= ≥ 2 experiments, ≥ 15 cells/ experiment, unpaired t-test vegetative 2% glucose vs 0% glucose low biotin p=.0006) All error bars are s.e.m.

**Supplementary Figure 11.**
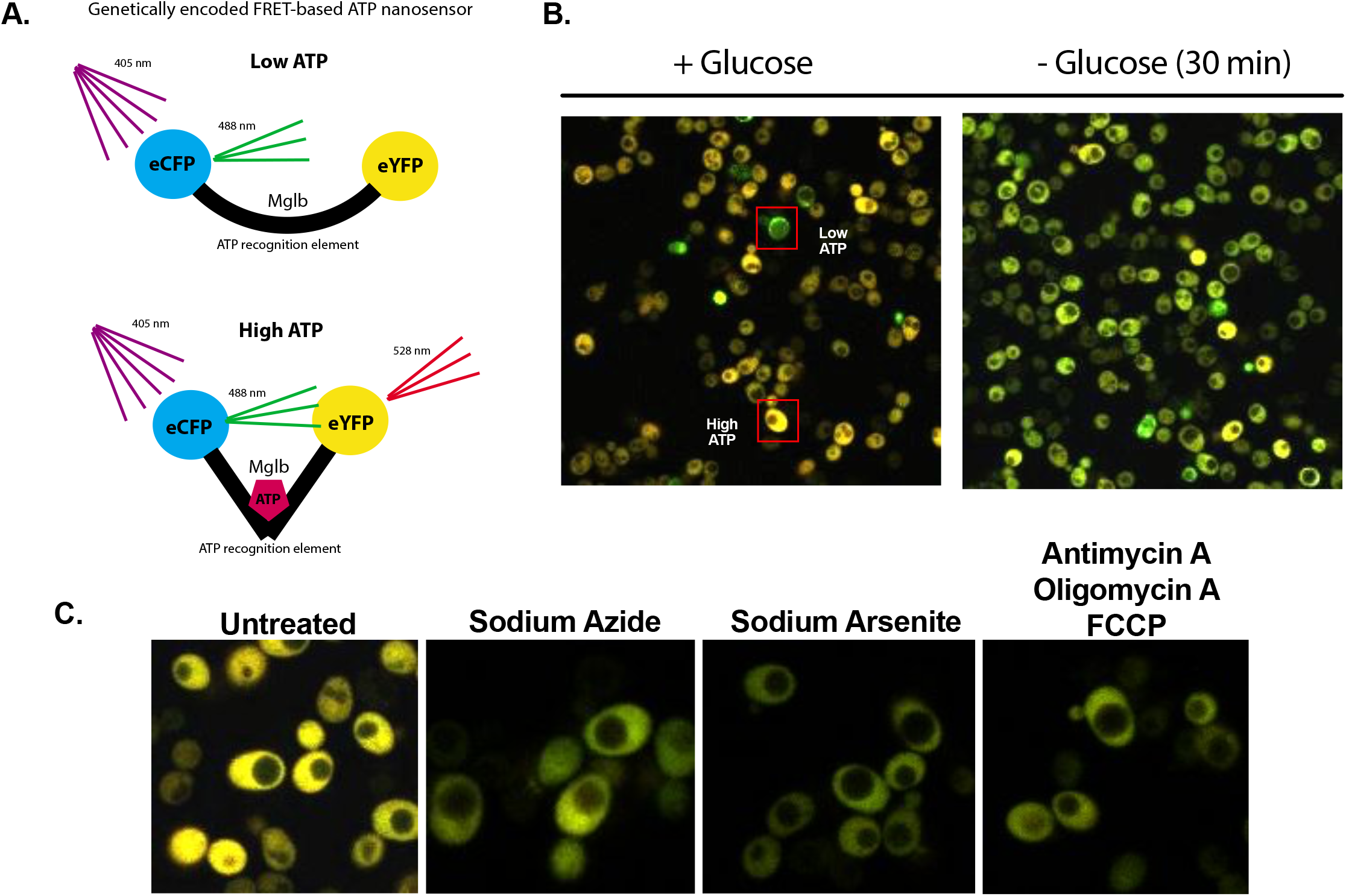
A ratiometric FRET-based sensor reports on relative ATP levels at single cell resolution. (**A**) Schematic representation of FRET based ATP sensor^33^. (**B**) Cells expressing the ATP sensor on a cen/ars plasmid in glucose replete (2%) or glucose starved (0%) conditions. Boxes highlight high and low ATP cells in the same culture. (**C**) Cells expressing the ATP sensor with the indicated drug treatments.

